# Recent and rapid anthropogenic habitat fragmentation increases extinction risk for freshwater biodiversity

**DOI:** 10.1101/2020.02.04.934729

**Authors:** Chris J. Brauer, Luciano B. Beheregaray

## Abstract

Anthropogenic habitat fragmentation is often implicated as driving the current global extinction crisis, particularly in freshwater ecosystems. The genetic signal of recent population isolation can however be confounded by the complex spatial arrangement of dendritic river systems. Consequently, many populations may presently be managed separately based on an incorrect assumption that they have evolved in isolation. Integrating landscape genomics data with models of connectivity that account for landscape structure, we show that the cumulative effects of multiple in-stream barriers have contributed to the recent decline of a freshwater fish from the Murray-Darling Basin, Australia. In addition, individual-based eco-evolutionary simulations further demonstrate that contemporary inferences about population isolation are consistent with the 160-year time frame since construction of in-stream barriers began in the region. Our findings suggest that the impact of very recent fragmentation may be often underestimated for freshwater biodiversity. We argue that proactive conservation measures to reconnect many riverine populations are urgently needed.

## Introduction

We are now confronted by the sixth global mass extinction with the current rate of species losses far exceeding pre-anthropogenic background estimates (Barnosky *et al.* 2011). This crisis is particularly severe in freshwater ecosystems, which have shown declines of biodiversity greater than for either terrestrial or marine ecosystems (Darwall *et al.* 2018). Habitat loss and fragmentation are key factors leading to the genetic and demographic decline of populations that together threaten species persistence (Fischer & Lindenmayer 2007). Over the last century, close to one million large dams and many millions of smaller in-stream barriers have been constructed globally (Jackson *et al.* 2001; Liermann *et al.* 2012). These barriers have had devastating ecological consequences by preventing or restricting connectivity among populations, leading to higher rates of genetic drift and inbreeding. This, in turn, can lead to lower fitness due to inbreeding depression and reduced evolutionary potential due to loss of genetic diversity (Frankham 2005; Keyghobadi 2007). Additionally, small populations become more vulnerable to extirpation due to stochastic demographic events (Lande 1993) and, when this occurs on a regional scale, species extinctions are the inevitable result (Hanski 1998).

Landscape genetics provides a way to identify how human activities threaten the persistence of wild populations (Manel & Holderegger 2013). The time lag between environmental change and any detectable genetic signal resulting from this change can however make it very difficult to disentangle the effects of historical from contemporary processes (Landguth *et al.* 2010). This is particularly the case for naturally structured populations such as those found in dendritic river networks (Coleman *et al.* 2018). The progression from landscape genetics to landscape genomics has increased both the spatial and temporal resolutions at which evolutionary processes can be examined, offering a more powerful framework with which to quantify the effects of very recent disturbance on populations (Allendorf *et al.* 2010; Grummer *et al.* 2019). Previous landscape genetics studies investigating the impact of in-stream barriers have often focused on larger, migratory species or assessed only one, or a few large barriers (Faulks *et al.* 2011; Gouskov *et al.* 2016; Torterotot *et al.* 2014). Small-bodied, but ecologically important species often receive relatively little attention from conservation managers (Olden *et al.* 2007; Saddlier *et al.* 2013) and efforts to improve fish passage and connectivity are often ineffective for these fishes (Harris *et al.* 2017). The cumulative impact of many smaller in-stream barriers (e.g. weirs, farm dams and road crossings) for small-bodied and non-migratory fishes at a basin-wide scale has been the subject of less research (Coleman *et al.* 2018; Diebel *et al.* 2015).

In this landscape genomics study, we examine the effects of recent habitat fragmentation for the southern pygmy perch (*Nannoperca australis*), a threatened small-bodied fish (<80mm) that recently experienced major demographic declines and local extinctions across the Murray-Darling Basin (MDB), Australia (Brauer *et al.* 2016; Cole *et al.* 2016; Hammer *et al.* 2013). This species is typical of many native small-bodied fishes in the region and offers a conservative model for guiding broader conservation strategies, as the impacts of fragmentation are likely to be more pronounced for larger, migratory species. The MDB has very few natural in-stream barriers, but it has been heavily modified with more than 10,000 dams, weirs, road crossings, levees and barrages constructed since the late 1850s when European settlement of this region began (Baumgartner *et al.* 2014). As such, the MDB provides a unique opportunity to examine the consequences of recent habitat fragmentation without the confounding influence of prolonged human disturbance over hundreds of years as is common to many northern hemisphere river basins (e.g. Hansen *et al.* 2014). Environmental factors, including human disturbance are known to influence genetic diversity for *N. australis* (Brauer *et al.* 2016; Cole *et al.* 2016), however little is known about the specific role that widespread habitat fragmentation has played in the species recent and rapid decline. We hypothesize that, after accounting for historical patterns of genetic structure, genetic differentiation among demes should increase with the number of in-stream barriers separating them. We also predict that populations most isolated by fragmentation would exhibit reduced effective population size (*N*_e_) and lower levels of genetic diversity.

Additionally, we used forward genetic simulations to investigate whether high contemporary levels of genetic differentiation could have arisen in the relatively short time since the construction of in-stream barriers began in the MDB. Our results demonstrate that recent anthropogenic habitat fragmentation has contributed to the loss of genetic diversity and population isolation observed. They also suggest that proactive conservation measures to restore connectivity (e.g. environmental flows, habitat restoration) and increase evolutionary potential (e.g. genetic rescue) are urgently required for this, and potentially many other poorly dispersing aquatic species.

## Methods

### Sampling, ddRAD genotyping and SNP filtering

A total of 263 individuals were sampled from 25 locations, encompassing 13 catchments across the entire current MDB distribution of *N. australis* (Figure 1; Table 1). Fish were ethically euthanized using clove oil, frozen in liquid nitrogen in the field, and stored at −70°C in the Australian Biological Tissues Collection at the South Australian Museum, Adelaide.

**Table 1.** Sample size (N), expected heterozygosity (*H*_E_), population-specific *F*_ST_ (Weir & Hill 2002) and effective population size estimates (*N*_e_). Lowland wetland sites referred to as Lower Murray in the text are indicated in bold. *MID and MUN samples combined for *N*_e_ estimation.

| Catchment | Site | N | $H_E$ | $F_{ST}$ | $N_e$ (95% CI) |
| --- | --- | --- | --- | --- | --- |
| **Tookayerta (TOO)** | **TBA** | **7** | **0.227** | **0.059** | ∞ |
| **Lower Lakes (LMR)** | **ALE** | **10** | **0.263** | **0.066** | **198.6 (158.6–264.9)** |
|  | **MID** | **7** | **0.262** | **0.092** | **190.9 (163.3–229.4)\*** |
|  | **MUN** | **6** | **0.260** | **0.034** |  |
| Angas (ANG) | MCM | 9 | 0.097 | 0.555 | 76.3 (61.0–101.3) |
| Avoca (AVO) | MIC | 11 | 0.114 | 0.409 | 13.7 (13.2–14.4) |
| Campaspe (CAM) | JHA | 12 | 0.091 | 0.364 | 393.8 (184.0–∞) |
| Upper Goulburn (UGO) | MER | 17 | 0.075 | 0.467 | 70.4 (61.4–82.2) |
|  | TRA | 10 | 0.075 | 0.433 | 50.7 (41.2–65.3) |
| | YEA | 8 | 0.087 | 0.364 | 260.4 (111.1–∞) |
| Lower Goulburn (LGO) | PRA | 9 | 0.243 | 0.179 | 114.9 (98.4–137.9) |
|  | SEV | 11 | 0.218 | 0.119 | 54.8 (50.8–59.4) |
| Broken (BRO) | BEN | 10 | 0.236 | 0.159 | 117.2 (101.7–138.2) |
|  | SAM | 10 | 0.234 | 0.188 | 124.7 (108.0–147.2) |
|  | LIM | 18 | 0.118 | 0.337 | 99.1 (88.5–112.5) |
| Ovens (OVE) | KIN | 16 | 0.104 | 0.297 | 69.9 (62.1–79.8) |
| | HAP | 9 | 0.114 | 0.369 | ∞ |
|  | MEA | 8 | 0.158 | 0.245 | 53.4 (45.7–64) |
| Kiewa (KIE) | GAP | 12 | 0.168 | 0.305 | 122.5 (105.3–146.2) |
| Albury (ALB) | ALB | 12 | 0.226 | 0.299 | 305.4 (241.8–413.4) |
| Mitta Mitta (MIT) | SPR | 10 | 0.152 | 0.262 | 98.1 (80.5–125) |
|  | GLE | 10 | 0.143 | 0.408 | 51.1 (46.1–57.2) |
|  | TAL | 7 | 0.164 | 0.479 | 31.9 (29.1–35.2) |
| Upper Murray (COP) | COP | 16 | 0.133 | 0.297 | 118.7 (102.2–141.1) |
| Lachlan (LAC) | LRT | 8 | 0.057 | 0.672 | 18.1 (15.3–21.8) |

**Figure 1.**
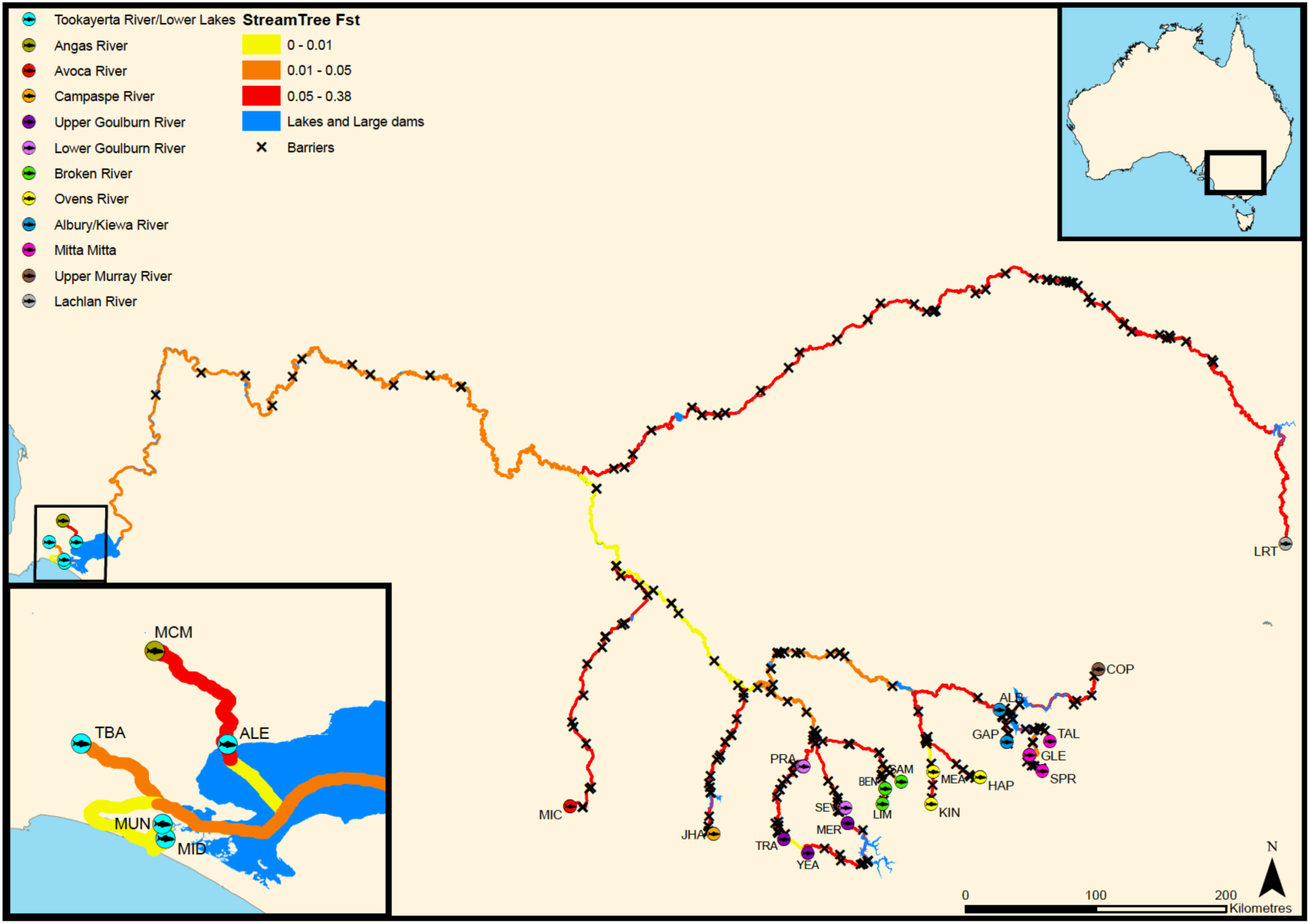
*Nannoperca australis* sampling locations in the Murray-Darling Basin (MDB). Stream sections are colour coded according to *F*_ST_ estimated using the *StreamTree* model (Kalinowski *et al.* 2008). Cross markers represent the location of artificial in-stream barriers.

DNA was extracted using the Qiagen DNeasy Blood and Tissue Kit according to the manufacturers protocol. DNA integrity and purity were assessed using gel electrophoresis and a NanoDrop 1000 spectrophotometer (Thermo Scientific), respectively. Sequencing libraries were prepared in-house based on a double-digest restriction-site associated DNA (ddRAD) library protocol (Peterson *et al.* 2012). Samples were multiplexed with 48 samples per Illumina HiSeq2000 lane and sequenced as paired-end, 100-bp reads. Raw sequences were demultiplexed using the *process_radtags* module of *Stacks* v.1.04 (Catchen *et al.* 2011) before using dDocent v.1.2 (Puritz *et al.* 2014) for *de novo* reference catalogue assembly and genotyping. The data was then filtered to retain only variants present in at least 70% of individuals and in 70% of populations, retaining only one biallelic SNP per locus with a minimum minor allele frequency of 0.05.

Population structure and other demographic parameters such as effective population size should be assessed using neutral loci (Allendorf *et al.* 2010; Luikart *et al.* 2003). To define a putatively neutral dataset, *F*_ST_ outlier loci were detected using a Bayesian approach with BayeScan v.2.1 (Foll & Gaggiotti 2008), and the coalescent-based FDIST method (Beaumont & Nichols 1996) in Arlequin v.3.5 (Excoffier & Lischer 2010). BayeScan was run for 100,000 iterations using prior odds of 10,000. Loci different from zero with a q-value <0.1 were considered outliers. Arlequin was run specifying the hierarchical island model with 50,000 simulations of 100 demes for each of 13 populations (based on the 13 separate catchments sampled). Loci outside the neutral distribution at a false discovery rate (FDR) of 10% were considered outliers. Loci detected as outliers by either BayeScan or Arlequin were filtered. The remaining SNPs were examined for departure from expectations of Hardy-Weinberg equilibrium (HWE) using GenoDive 2.0b27 (Meirmans & Van Tienderen 2004). Finally, loci out of HWE at a FDR of 10% in more than 50% of populations were removed. Detailed information concerning library preparation and bioinformatics are described in Appendix S1.

### Population structure

Pairwise *F*_ST_ (Weir & Cockerham 1984) was estimated among sampling sites using GenoDive (Meirmans & Van Tienderen 2004) with significance assessed using 10,000 permutations. Bayesian clustering analysis of individual genotypes was then performed using fastStructure (Raj *et al.* 2014). Ten independent runs for each value of *K* (1-25) were completed to ensure consistency and the most likely *K* was assessed by comparing the model complexity that maximised marginal likelihood across replicate runs.

### Anthropogenic isolation of populations

If anthropogenic habitat fragmentation has affected population connectivity and dispersal, we should expect genetic differentiation to increase in response to the number of in-stream barriers separating populations. To determine if local characteristics of the stream network (i.e. in-stream barriers and other local scale landscape heterogeneity) better explain population differentiation than isolation by distance (IBD), we used the StreamTree model of Kalinowski et al. (2008). Genetic distances among populations were modelled as the sum of all pairwise genetic distances that mapped to each section of the stream network. This provides a distance measure that is independent of the length of each stream section and identifies the reaches that contribute most to restricting gene flow (e.g. due to dendritic structure, in-stream barriers or other local landscape effects). Model fit was assessed by plotting the StreamTree fitted distance against observed *F*_ST_ and calculating the regression coefficient of determination (*R*^2^). This model was then compared with a model of IBD calculated using multiple matrix regression with randomisation (MMRR) following the method of Wang (2013). Pairwise population distances along the river network were calculated with ArcMap v.10.2 (ESRI 2012). Model significance for the MMRR was assessed using 10,000 random permutations.

In dendritic river systems, hierarchical network structure and spatial hydroclimatic variation can also drive patterns of genetic diversity of stream-dwelling organisms (Fourcade *et al.* 2013; Hughes *et al.* 2009; Morrissey & de Kerckhove 2009; Thomaz *et al.* 2016). To evaluate the relative contributions of anthropogenic habitat fragmentation, natural stream hierarchy and environmental variation we again used MMRR. In addition to IBD, we used distance matrices calculated for the number of in-stream barriers, catchment membership, and a range of environmental variables. The number of in-stream barriers separating sites was determined using spatial data from the Murray-Darling Basin Weir Information System (Murray–Darling Basin Authority 2013). To account for the effect of dendritic stream hierarchy, a binary model matrix describing catchment membership was constructed such that pairwise comparisons of sites from within the same catchment were assigned a value of zero, and comparisons among catchments were scored as one. Finally, a subset of 40 hydroclimatic variables were obtained from the Australian hydrological geospatial fabric (Geoscience Australia 2011; Stein *et al.* 2014). These were assigned to one of five categories describing variation in temperature, precipitation, flow regime, human disturbance and topography. Variance inflation factor (VIF) analysis was then used to exclude highly correlated variables using a VIF threshold of 10 (Dyer *et al.* 2010). The remaining variables were reduced to principal components (PCs) using the dudi.pca function in the ADE4 R package (Dray *et al.* 2016) and Euclidean distance matrices were constructed based on the PCs with eigenvalues >1 (Yeomans & Golder 1982) retained for each category. All distance matrices were z-transformed to facilitate direct comparison of partial regression coefficients (Schielzeth 2010). Each variable was initially tested in an independent univariate MMRR before significant factors were combined in a multivariate MMRR model with 10,000 random permutations used to assess significance.

### Habitat fragmentation, genetic diversity and population size

To test the hypothesis that the most isolated populations exhibit reduced genetic diversity we examined the relationship between population-specific *F*_ST_ and expected heterozygosity (*H*_E_). Population-specific *F*_ST_ was estimated for each sampling site using the method of Weir and Hill (2002) and *H*_E_ was calculated using Genodive.

Effective population size was estimated using the linkage disequilibrium (LD) estimator implemented in NeEstimator 2.01 (Do *et al.* 2014). This method is based on the assumption that LD at independently segregating loci in a finite population is a function of genetic drift, and performs particularly well with a large number of loci and where population sizes are expected to be small (Waples & Do 2010). In the absence of significant *F*_ST_ (Table S1) Lower Murray sites MID and MUN were considered one population, and these samples were combined for the *N*_e_ estimates. NeEstimator was run assuming random mating and using a *P*_crit_ value of 0.075 following guidelines for small sample sizes suggested by Waples and Do (2010).

### Eco-evolutionary simulations

Simulation studies are becoming an increasingly important part of landscape genomics as a wide range of parameters can be explored for key evolutionary processes such as gene flow, genetic drift, mutation and selection (Hoban *et al.* 2012). In this case, we used simulations to examine whether levels of contemporary population isolation are consistent with having evolved during the time since barrier construction began in the MDB. We simulated three metapopulation sizes (*N*_e_=1000, *N*_e_=500 and *N*_e_=100) using SLiM 3.1 (Haller & Messer 2018). Each simulation was based on a 1D stepping stone population model assuming equal *N*_e_ for each sub-population while maintaining a constant metapopulation size to simulate a concurrent increase in the number of barriers, and reduction in habitat patch size. Each simulation consisted of four 100Kb genomic elements and assumed a constant mutation rate of 10^−7^ and recombination rate of 10^−8^. Each simulation was first run for a burn-in phase of 20,000 generations with a migration rate of 0.5 between adjacent sub-populations to generate diversity and allow the system to reach migration–drift equilibrium with *F*_ST_=∼0. Although this almost certainly underestimates historical population structure before anthropogenic disturbance, this figure provides a conservative approach by maximizing the number of generations required to evolve current levels of differentiation. Following the burn-in, the construction of barriers was simulated by setting the migration rate among demes to zero for 300 generations. Nine models with an increasing number of demes (2-10) were simulated for each metapopulation size to examine the effect of increasing levels of fragmentation (Figure S1-S3), and 100 replicate runs of each scenario were completed. The --weir-fst-pop command of VCFtools (Danecek *et al.* 2011) was used to calculate *F*_ST_ for each replicate. To estimate the time required to reach current levels of observed population differentiation, assuming a generation time of one year (Humphries 1995), the number of generations (mean of the 100 replicates) needed to achieve *F*_ST_=0.2 (mean contemporary *F*_ST_ within upper Murray catchments=0.196; Table S1) was plotted against the number of fragments for each scenario for the three metapopulation models. Scripts used to perform the simulations and analyses are available on Dryad: TBA.

## Results

### Sampling, ddRAD genotyping and SNP filtering

Following demultiplexing, 1,602,903,910 forward and reverse sequencing reads were recovered. A total of 2,589,251 variant sites were genotyped with dDocent, and after filtering 5,162 high quality SNPs were retained. We removed 873 unique *F*_ST_ outlier loci identified by BayeScan and Arlequin, along with a further 846 loci found to be outside HWE expectations in >50% of populations. This resulted in a final, putatively neutral dataset of 3,443 SNPs for the 263 individuals.

### Population structure

High levels of population genetic structure were evident between most demes of *N. australis*, with pairwise comparisons of *F*_ST_ among sampling sites ranging from 0-0.79 (global *F*_ST_ =0.48). All pairwise *F*_ST_ estimates were significant (*P*<0.003) except between immediately adjacent lower MDB sites MID and MUN (*F*_ST_= −0.002, *P*=0.66) (Table S1). Results from fastStructure indicated population boundaries are strongly correlated with natural riverine catchment structure, with K=13 identified as the most likely number of populations (Figure S4). This is consistent with a previous microsatellite study based on a larger sample (578 individuals; 45 localities) that inferred that, until the recent European settlement in the MDB, well-connected metapopulations of *N. australis* existed within its catchments (Cole *et al.* 2016).

### Anthropogenic isolation of populations

The StreamTree model was used to identify parts of the stream network that contribute more to *F*_ST_ (e.g. restricted dispersal due to barriers or other local environmental conditions). Results indicated that local characteristics of the stream network better explain *F*_ST_ than the null hypothesis of IBD (i.e. the resistance to dispersal for any given stream section is determined by its length). Figure 1 provides a visual representation of the relationship between StreamTree fitted distance and the density of artificial in-stream barriers, with stream sections colour coded according to *F*_ST_ as estimated by the model (yellow represents a modeled local *F*_ST_ range of 0-0.01, orange: 0.01-0.05 and red: 0.05-0.38) and the location of barriers marked with **X**. The StreamTree model was a good fit for the data and was significantly related to observed *F*_ST_ (*R*^2^=0.947, β=0.986 [0.959- 1.012 95%CI], *P*<2×10^−16^) (Figure 2a), whereas IBD was not significant (*R*^2^=0.0139, β=0.108 [0.004- 0.212 95%CI], *P*=0.343) (Figure 2b). Although there was significant IBD within catchment groups (i.e. the first cluster in Figure 2b, *R*^2^=0.730, β=0.0016 [0.001- 0.002 95%CI], *P*=6.54×10^−8^), IBD was not significant in models across the whole basin, in contrast to models of stream hierarchy and barriers (see below). In addition, even when comparisons were limited to sites within-catchments, the number of barriers still provided a better model than IBD (*R*_2_=0.81 vs. 0.73, respectively; Figure S5).

**Figure 2.**
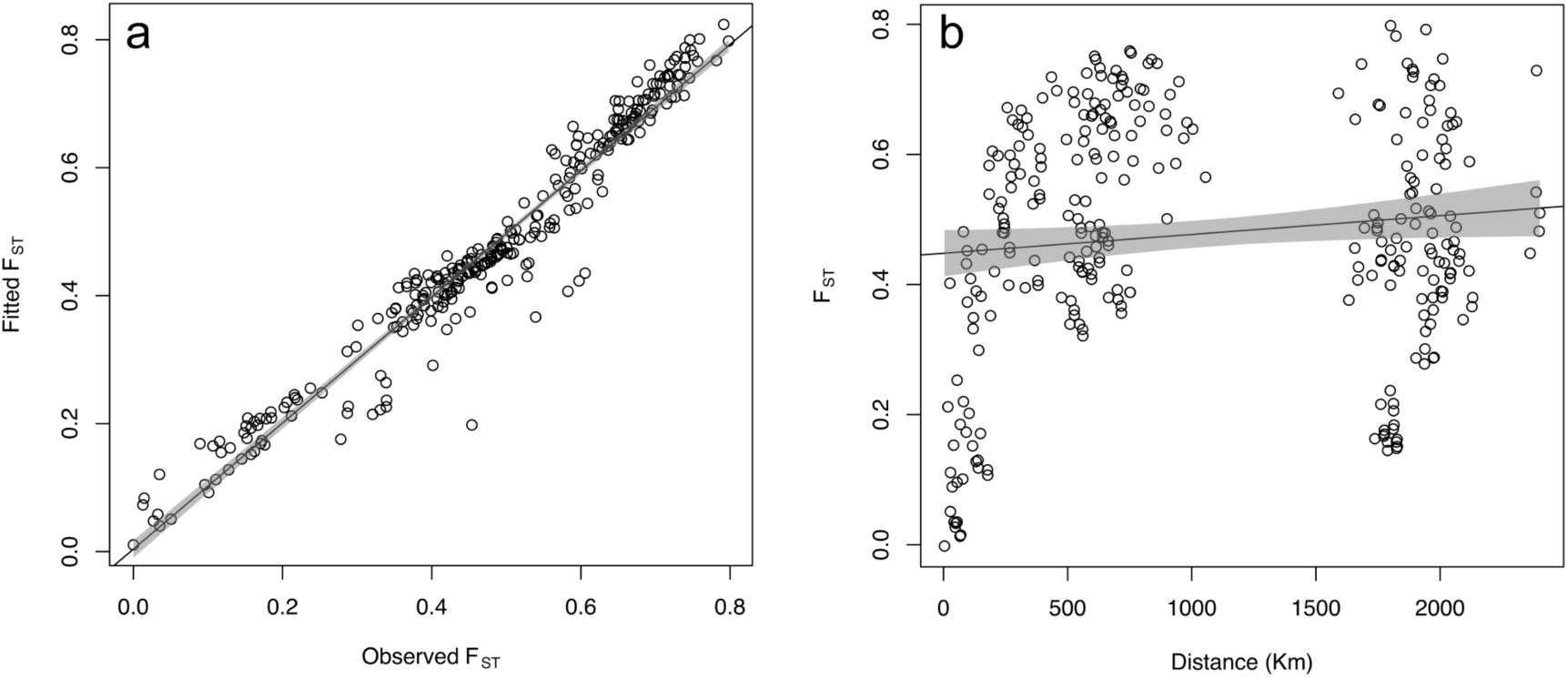
Plots of a) *StreamTree* analyses and b) isolation by distance (IBD) for *Nannoperca australis* in the MDB. The *StreamTree* plot compares fitted *F*_ST_ based on the *StreamTree* model with observed pairwise *F*_ST_ values (*R*^2^=0.947, β=0.986 [0.959- 1.012 95%CI], *P*<2×10^−16^). The IBD plot depicts the relationship between pairwise *F*_ST_ and riverine distance between sampling sites (*R*^2^=0.0139, β=0.108 [0.004- 0.212 95%CI], *P*=0.343). Shaded area represents the 95% confidence interval.

Following VIF analyses, 19 environmental variables from the five categories were retained. The first two PCs for temperature, flow and topographic variables scored eigenvalues >1 while only one component each for the precipitation and human disturbance PCAs scored an eigenvalue >1, so individual variables rather than PCs for these categories were retained. This resulted in a final list of six hydroclimatic PCs and five individual precipitation and disturbance variables (Table S2).

Assessment of the relative influence of anthropogenic habitat fragmentation, natural stream hierarchy and environmental heterogeneity indicated that population structure is driven by a combination of the effects of stream network hierarchy and the number of in-stream barriers. Univariate regressions revealed catchment membership (*R*^2^=0.170, β=0.449 [0.336- 0.562 95%CI], *P*<0.0001) and the number of in-stream barriers separating sites (*R*^2^=0.322, β=0.548 [0.458- 0.639 95%CI], *P*<0.0001) were both good predictors of population differentiation, while there was no evidence for isolation by environment (Table 2). Including both significant predictors (catchment membership and number of barriers) in a multivariate model improved model fit with catchment membership, and the number of barriers each accounting for 61% and 39% of the explained variation, respectively (*R*^2^=0.358, β_catchment_=0.725 [0.374- 1.076 95%CI], β_barriers_=0.462 [0.365- 0.560 95%CI], *P*<0.0001) (Figure 3; Table 2).

**Table 2.** Results of multiple matrix regression with randomisation (MMRR) tests for the relationship between pairwise genetic distance (*F*_ST_) and geographic distance, catchment membership, number of in-stream barriers and environmental distances. Pairwise environmental distances between each site were calculated as Euclidean distance for each environmental variable and principal component (PC) described in Brauer *et al*. (2016). *P*-values <0.0001 are indicated in bold.

| Model | Variable | Coefficient | 95%CI | *P*-value | $R^2$ | Model *P*-value |
| --- | --- | --- | --- | --- | --- | --- |
|  | Distance | 0.108 | 0.004- 0.213 | 0.3340 | 0.014 |  |
|  | Catchment | 0.449 | 0.336- 0.562 | **0.0001** | 0.170 |  |
|  | Barriers | 0.548 | 0.458- 0.639 | **0.0001** | 0.322 |  |
|  | TempPC1 | -0.130 | -0.233- -0.028 | 0.2465 | 0.021 |  |
|  | TempPC2 | 0.180 | 0.077- 0.282 | 0.1443 | 0.039 |  |
|  | CATCOLDQRAIN | 0.098 | -0.007- 0.202 | 0.3813 | 0.011 |  |
|  | CATDRYQRAIN | -0.061 | -0.170- 0.043 | 0.5515 | 0.004 |  |
|  | STRWETQRAIN | -0.058 | -0.162- 0.046 | 0.5496 | 0.004 |  |
|  | FlowPC1 | -0.053 | -0.158- 0.051 | 0.6698 | 0.003 |  |
|  | FlowPC2 | -0.125 | -0.227- -0.023 | 0.3520 | 0.019 |  |
|  | CDI | 0.037 | -0.068- 0.142 | 0.6571 | 0.002 |  |
|  | FRDI | -0.087 | -0.190- 0.015 | 0.4603 | 0.009 |  |
|  | TopoPC1 | -0.121 | -0.225- -0.017 | 0.2368 | 0.017 |  |
|  | TopoPC2 | 0.021 | -0.083- 0.125 | 0.8644 | 0.001 |  |
| Catchment+Barriers |  |  |  |  | 0.358 | **0.0001** |
|  | Catchment | 0.725 | 0.374- 1.076 | **0.0045** |  |  |
|  | Barriers | 0.462 | 0.365- 0.560 | **0.0001** |  |  |

**Figure 3.**
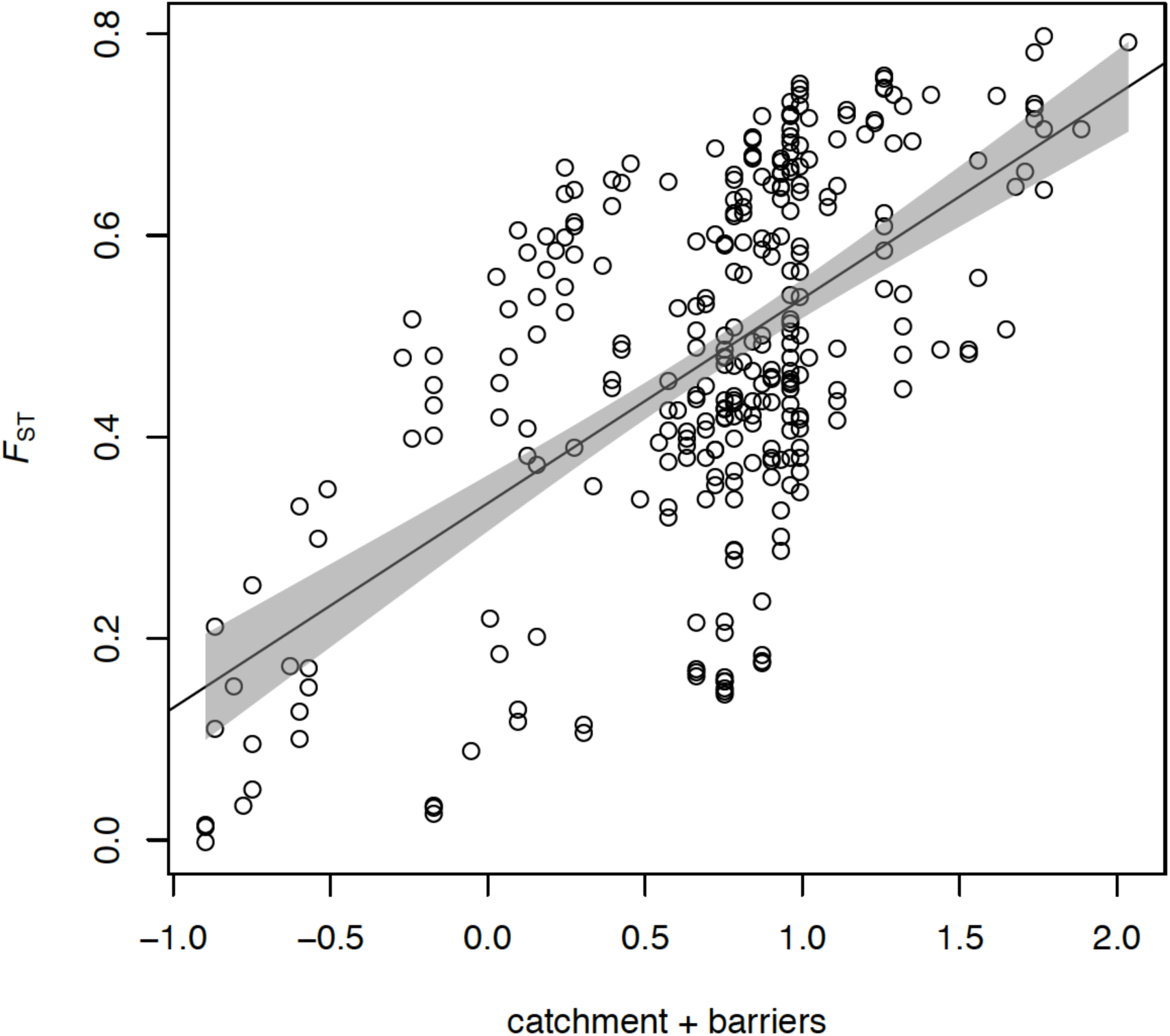
Multiple matrix regression with randomisation (MMRR) plot for the combined effects of natural stream hierarchy (model matrix of catchment membership) and number of barriers on *F*_ST_ (*R*^2^=0.358, β_catchment_=0.725 [0.374- 1.076 95%CI], β_barriers_=0.462 [0.365- 0.560 95%CI], *P*<0.0001). Shaded area represents the 95% confidence interval.

### Habitat fragmentation, genetic diversity and population size

Genetic variation varied across the MDB with an average *H*_*E*_ of 0.161 (0.057-0.263). There was a sharp contrast between regions with average *H*_*E*_ of 0.253 for sites in the more connected Lower Murray wetlands, compared to 0.143 for sites in the highly fragmented upper reaches (Table 1). A strong negative relationship between population-specific *F*_ST_ and *H*_*E*_ was also evident (*R*^2^=0.737, β=-2.05 [-2.58- -1.52 95%CI], *P*<1×10^−7^) with the most isolated populations also harbouring the least genetic variation (Figure S6; Table 1). Effective population size estimates were generally low, averaging 194.75 for Lower Murray sites and 112.26 for sites in the upper reaches, with many of the latter <100 (Table 1).

### Eco-evolutionary simulations

The simulations demonstrated that contemporary population differentiation among sites within catchments (mean within headwater catchments *F*_ST_=0.196) could have evolved from a more connected system within the time since the construction of in-stream barriers began ∼160 generations ago (Figure 4; Table S3; Appendix S3-S5). For metapopulations with an *N*_e_ of 1000, *F*_ST_ approached 0.2 in less than 160 generations with only three barriers fragmenting the population. Models assuming *N*_e_=500 and *N*_e_=100 indicated that substantially fewer generations following fragmentation were required to reach contemporary levels of *F*_ST_. At *N*_e_=500, *F*_ST_ =0.2 occurred after 124 generations with one barrier and after just 19 generations with nine barriers (Figure 4; Table S3). For smaller populations of *N*_e_=100 contemporary levels of differentiation evolved within 24 generations with just one barrier (Figure 4; Table S3).

**Figure 4.**
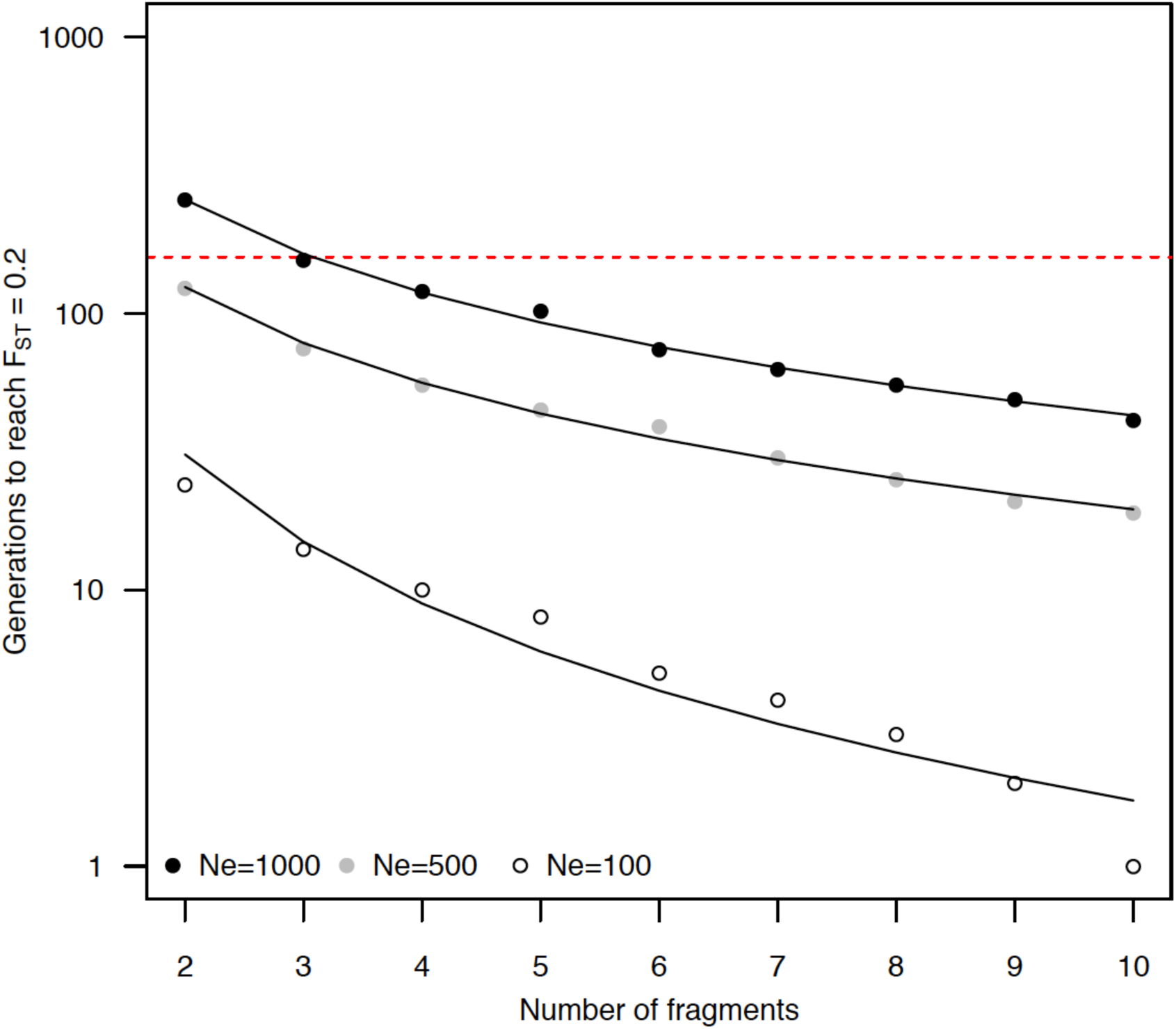
Number of generations (log scale) for global *F*_ST_ to reach 0.2 with increasing levels of habitat fragmentation for simulated *N. australis* metapopulations of *N*_e_=1000, *N*_e_ =500 and *N*_e_ =100. Simulations were based on a stepping stone model assuming equal *N*_e_ for each sub-population and were allowed to run for 20,000 generations with a migration rate of 0.5 between adjacent demes before 300 generations with no migration. Red dashed line indicates the approximate number of generations since construction of in-stream barriers began in the MDB (160 generations).

## Discussion

Habitat fragmentation is a key process implicated in the current and unprecedented worldwide loss of freshwater biodiversity (Fischer & Lindenmayer 2007). Determining the contribution of recent human activities to the decline of riverine species is however challenging, as the genetic signal of recent disturbance can be confounded by historical patterns of dispersal shaped by hydrological network structure (Brauer *et al.* 2018; Coleman *et al.* 2018; Landguth *et al.* 2010). For *N. australis*, populations most isolated by recent habitat fragmentation also exhibited reduced genetic diversity and increased population differentiation, and this signal remained strong after accounting for the historical effects of dendritic stream hierarchy. Contemporary *F*_ST_ estimates were within the expected range obtained by simulating the recent construction of in-stream barriers in the MDB, supporting the hypothesis that anthropogenic habitat fragmentation has impacted populations since European settlement of the region. Previous work based on coalescent analyses of microsatellite DNA datasets has demonstrated that historical population sizes of *N. australis* were much larger before European settlement (Attard *et al.* 2016), and that populations across the MDB were also more connected until that time (Cole *et al.* 2016). These studies support our findings and the hypothesis that the low genetic diversity, small *N*_e_ and high *F*_ST_ observed for contemporary populations likely reflects the combined impact of both historical and recent processes, rather than being due solely to natural demographic variability over longer evolutionary time scales. In addition, several populations sampled for this study have subsequently suffered local extirpation during prolonged drought, and the small size of most remnant populations indicate they are at high risk of extinction.

Since European settlement, hydrology in the MDB has been increasingly modified due to urbanisation and irrigation (Leblanc *et al.* 2012). These changes have included the construction of thousands of barriers to fish passage across the basin (Baumgartner *et al.* 2014) and it is now considered one of Australia’s most fragmented and degraded ecosystems (Davies *et al.* 2010; Kingsford 2000). The focus of most barrier mitigation actions in the MDB to date has been on restoring passage across larger dams along the main river channel (Barrett & Mallen-Cooper 2006). Although some fishways have been designed to facilitate movement of smaller fish, they have predominantly targeted large-bodied, highly mobile species (Baumgartner *et al.* 2014). Furthermore, the spatial scale of dispersal for many small-bodied MDB fishes often restricts their movements to headwater streams and wetlands away from the main channel (Harris *et al.* 2017). Habitat loss and fragmentation associated with the thousands of smaller barriers in headwater streams have therefore likely contributed to the widespread decline of many smaller and more sedentary MDB fishes, including *N. australis* (Brauer *et al.* 2018; Cole *et al.* 2016; Hammer *et al.* 2013; Huey *et al.* 2017). It is perhaps surprising then, that there have been relatively few studies explicitly testing the genetic effects of anthropogenic fragmentation on small-bodied fishes in the MDB. One recent example in the neighbouring Yarra River catchment however combined a large empirical dataset with spatially explicit simulations to examine the role of artificial barriers in driving local-scale patterns of genetic variation for river blackfish (*Gadopsis marmoratus*), a small and sedentary species also found in the MDB (Coleman *et al.* 2018). Based on eight microsatellite loci, genetic diversity was found to be lower for populations above barriers in small streams, with several isolated populations also exhibiting signs of inbreeding. In addition, their simulations demonstrated that power to detect recent impacts of barriers could be improved by increasing the number of loci used, highlighting the benefit of modern genomic data for conservation genetics.

An unprecedented severe and prolonged drought between 1997 and 2010 caused catastrophic loss of habitat and local extirpation for some *N. australis* populations, particularly in the lower Murray (Hammer *et al.* 2013; Wedderburn *et al.* 2012). In response an emergency conservation-breeding and restoration program was implemented in the lower MDB (Attard *et al.* 2016; Hammer *et al.* 2013), and additional breeding and translocations among several headwater populations have been initiated (P. Rose, *personal communication*). As the impacts of climate change intensify, proactive conservation management interventions such as those already underway for *N. australis*, will be increasingly considered for other species inhabiting the MDB and fragmented freshwater ecosystems elsewhere in the world. Indeed, a recent study incorporating physiological and functional traits with species distribution models for 23 fish species predicted severe declines in taxonomic and functional diversity of MDB fish communities in the coming decades due to climate change (de Oliveira *et al.* 2019). Managing regulated river systems to provide environmental flows, habitat restoration and other measures to re-establish connectivity among habitat patches (e.g. installation of fishways) have the potential to address some impacts and should continue to be priorities for conservation and water management. Nonetheless, these long-term, landscape-scale measures are often constrained by competing interests related to political and socio-economic issues (Davis *et al.* 2015).

Additionally, many species may be already depleted to the point where improved environmental conditions alone will not be sufficient to facilitate recovery. In this case, genetic rescue offers a potential solution for a broad range of threatened taxa (Ralls *et al.* 2018; Whiteley *et al.* 2015). Despite strong evidence supporting the benefits of genetic rescue for fragmented populations however, conservation managers are often reluctant to adopt these measures (Frankham 2015). We suggest that the impacts of recent habitat fragmentation may have been underappreciated for many species, and that estimates of population structure solely attributed to historical evolutionary processes have potentially led to management frameworks that actually reinforce fragmentation and isolation at the expense of species-level genetic variation (sensu Coleman *et al.* 2013). There is also increasing evidence that natural selection can influence the evolutionary trajectory of small and fragmented populations (Brauer *et al.* 2017; Fraser 2017; Wood *et al.* 2016). Critically for conservation, this indicates that adaptive divergence of small populations can occur quickly following fragmentation (Brauer *et al.* 2017), and that even very recently isolated populations may harbor novel adaptive diversity. It is therefore important to build evolutionary resilience by facilitating genetic exchange among isolated populations to restore natural evolutionary processes and maintain species-level genetic variation, potentially valuable under a range of future selection regimes (Webster *et al.* 2017; Weeks *et al.* 2016).

There is a global biodiversity crisis unfolding in freshwater ecosystems with aquatic vertebrate populations declining by 80% over the last 50 years (Darwall *et al.* 2018). Restoring functional connectivity for whole aquatic communities across entire river basins via traditional mitigation approaches is simply not feasible within the time frame required to enable many currently threatened species to persist. There is also now strong empirical evidence that several long-established beliefs central to prevailing conservation practices are overly cautious, and that the current local-is-best approach increases the prospect of managing species to extinction (Frankham *et al.* 2017; Pavlova *et al.* 2017; Weeks *et al.* 2016). Given widespread fragmentation, habitat loss, and the ongoing global decline of freshwater biodiversity, a rapid paradigm shift is needed to empower conservation practitioners to take action before a threatened species demographic situation becomes critical. There are risks associated with any proactive management intervention such as translocation or genetic rescue. These risks however need to be weighed against the ever-increasing risk of doing nothing.

## Acknowledgements

We thank the many people who helped with fieldwork, especially Peter Unmack, Michael Hammer and Mark Adams for curating samples. Minami Sasaki and Jonathan Sandoval-Castillo provided valuable assistance with laboratory and bioinformatics, respectively. Collections were obtained under permits from various state fisheries agencies and research is under Flinders University Animal Welfare Committee approval E313. Financial support was provided by the Australian Research Council via a Future Fellowship project to Luciano Beheregaray (FT130101068) and the AJ & IM Naylon PhD scholarship to Chris Brauer.

## Data Archiving Statement

Reference sequences, SNP genotypes, sample coordinates and environmental data used in analyses are available on Dryad: TBA

## References

Allendorf FW, Hohenlohe PA, Luikart G (2010) Genomics and the future of conservation genetics. Nature Reviews: Genetics 11, 697–709.

Attard C, Möller L, Sasaki M, et al. (2016) A novel holistic framework for genetic-based captive-breeding and reintroduction programs. Conservation Biology 30, 1060–1069.

Barnosky AD, Matzke N, Tomiya S, et al. (2011) Has the Earth’s sixth mass extinction already arrived? Nature 471, 51–57.

Barrett J, Mallen-Cooper M (2006) The Murray River’s ‘Sea to Hume Dam’ fish passage program: Progress to date and lessons learned. Ecological Management & Restoration 7, 173–183.

Baumgartner L, Zampatti B, Jones M, Stuart I, Mallen-Cooper M (2014) Fish passage in the Murray-Darling Basin, Australia: Not just an upstream battle. Ecological Management & Restoration 15, 28–39.

Beaumont MA, Nichols RA (1996) Evaluating loci for use in the genetic analysis of population structure. Proceedings of the Royal Society of London. Series B: Biological Sciences 263, 1619–1626.

Brauer C, Unmack PJ, Smith S, Bernatchez L, Beheregaray LB (2018) On the roles of landscape heterogeneity and environmental variation in determining population genomic structure in a dendritic system. Molecular Ecology 27, 3484–3497.

Brauer CJ, Hammer M, Beheregaray L (2016) Riverscape genomics of a threatened fish across a hydroclimatically heterogeneous river basin. Molecular Ecology 25, 5093–5113.

Brauer CJ, Unmack PJ, Beheregaray LB (2017) Comparative ecological transcriptomics and the contribution of gene expression to the evolutionary potential of a threatened fish. Molecular Ecology 26, 6841–6856.

Catchen JM, Amores A, Hohenlohe P, Cresko W, Postlethwait JH (2011) Stacks: building and genotyping loci de novo from short-read sequences. G3: Genes, Genomes, Genetics 1, 171–182.

Cole T, Hammer M, Unmack P, et al. (2016) Range-wide fragmentation in a threatened fish associated with post-European settlement modification in the Murray-Darling Basin, Australia. Conservation Genetics 17, 1377–1391.

Coleman RA, Gauffre B, Pavlova A, et al. (2018) Artificial barriers prevent genetic recovery of small isolated populations of a low-mobility freshwater fish. Heredity 120, 515–532.

Coleman RA, Weeks AR, Hoffmann AA (2013) Balancing genetic uniqueness and genetic variation in determining conservation and translocation strategies: a comprehensive case study of threatened dwarf galaxias, *Galaxiella pusilla* (Mack) (Pisces: Galaxiidae). Molecular Ecology 22, 1820–1835.

Danecek P, Auton A, Abecasis G, et al. (2011) The variant call format and VCFtools. Bioinformatics 27, 2156–2158.

Darwall W, Bremerich V, De Wever A, et al. (2018) The Alliance for Freshwater Life: A global call to unite efforts for freshwater biodiversity science and conservation. Aquatic Conservation: Marine and Freshwater Ecosystems 28, 1015–1022.

Davies P, Harris J, Hillman T, Walker K (2010) The sustainable rivers audit: assessing river ecosystem health in the Murray–Darling Basin, Australia. Marine and Freshwater Research 61, 764–777.

Davis J, O’Grady AP, Dale A, et al. (2015) When trends intersect: the challenge of protecting freshwater ecosystems under multiple land use and hydrological intensification scenarios. Science of the Total Environment 534, 65–78.

de Oliveira AG, Bailly D, Cassemiro FA, et al. (2019) Coupling environment and physiology to predict effects of climate change on the taxonomic and functional diversity of fish assemblages in the Murray-Darling Basin, Australia. PLoS ONE 14.

Diebel M, Fedora M, Cogswell S, O’Hanley J (2015) Effects of road crossings on habitat connectivity for stream-resident fish. River Research and Applications 31, 1251–1261.

Do C, Waples RS, Peel D, et al. (2014) NeEstimator v2: re-implementation of software for the estimation of contemporary effective population size (*N*e) from genetic data. Molecular Ecology Resources 14, 209–214.

Dray S, Legendre P, Blanchet FG (2016) packfor: Forward Selection with permutation R package version 0.0-8/r136. https://R-Forge.R-project.org/projects/sedar/.

Dyer RJ, Nason JD, Garrick RC (2010) Landscape modelling of gene flow: improved power using conditional genetic distance derived from the topology of population networks. Molecular Ecology 19, 3746–3759.

ESRI (2012) ArcMap 10.1. Environmental Systems Research Institute, Inc., Redlands.

Excoffier L, Lischer HEL (2010) Arlequin suite ver 3.5: a new series of programs to perform population genetics analyses under Linux and Windows. Molecular Ecology Resources 10, 564–567.

Faulks LK, Gilligan DM, Beheregaray LB (2011) The role of anthropogenic vs. natural in-stream structures in determining connectivity and genetic diversity in an endangered freshwater fish, Macquarie perch (*Macquaria australasica*). Evolutionary Applications 4, 589–601.

Fischer J, Lindenmayer DB (2007) Landscape modification and habitat fragmentation: a synthesis. Global Ecology and Biogeography 16, 265–280.

Foll M, Gaggiotti O (2008) A genome-scan method to identify selected loci appropriate for both dominant and codominant markers: a bayesian perspective. Genetics 180, 977–993.

Fourcade Y, Chaput-Bardy A, Secondi J, Fleurant C, Lemaire C (2013) Is local selection so widespread in river organisms? Fractal geometry of river networks leads to high bias in outlier detection. Molecular Ecology 22, 2065–2073.

Frankham R (2005) Genetics and extinction. Biological Conservation 126, 131–140.

Frankham R (2015) Genetic rescue of small inbred populations: meta-analysis reveals large and consistent benefits of gene flow. Molecular Ecology 24, 2610–2618.

Frankham R, Ballou JD, Ralls K, et al. (2017) Genetic management of fragmented animal and plant populations Oxford University Press.

Fraser DJ (2017) Genetic diversity of small populations: Not always “doom and gloom”? Molecular Ecology 26, 6499–6501.

Geoscience Australia (2011) National surface water information. http://www.ga.gov.au/topographic-mapping/national-surface-water-information.html

Gouskov A, Reyes M, Wirthner-Bitterlin L, Vorburger C (2016) Fish population genetic structure shaped by hydroelectric power plants in the upper Rhine catchment. Evolutionary Applications 9, 394–408.

Grummer JA, Beheregaray LB, Bernatchez L, et al. (2019) Aquatic Landscape Genomics and Environmental Effects on Genetic Variation. Trends in Ecology & Evolution.

Haller BC, Messer PW (2018) SLiM 3: Forward genetic simulations beyond the Wright– Fisher model. Molecular Biology and Evolution.

Hammer MP, Bice CM, Hall A, et al. (2013) Freshwater fish conservation in the face of critical water shortages in the southern Murray–Darling Basin, Australia. Marine and Freshwater Research 64, 807–821.

Hansen MM, Limborg MT, Ferchaud A-L, Pujolar J-M (2014) The effects of Medieval dams on genetic divergence and demographic history in brown trout populations. BMC Evolutionary Biology 14, 1.

Hanski I (1998) Metapopulation dynamics. Nature 396, 41–49.

Harris JH, Kingsford RT, Peirson W, Baumgartner LJ (2017) Mitigating the effects of barriers to freshwater fish migrations: the Australian experience. Marine and Freshwater Research 68, 614–628.

Hoban S, Bertorelle G, Gaggiotti OE (2012) Computer simulations: tools for population and evolutionary genetics. Nature Reviews: Genetics 13, 110–122.

Huey JA, Balcombe SR, Real KM, Sternberg D, Hughes JM (2017) Genetic structure and effective population size of the most northern population of the Australian river blackfish, *Gadopsis marmoratus* (Richardson 1848): implications for long-term population viability. Freshwater Science 36, 113–123.

Hughes JM, Schmidt DJ, Finn DS (2009) Genes in streams: using DNA to understand the movement of freshwater fauna and their riverine habitat. Bioscience 59, 573–583.

Humphries P (1995) Life history, food and habitat of southern pygmy perch, *Nannoperca australis*, in the Macquarie River, Tasmania. Marine and Freshwater Research 46, 1159–1169.

Jackson RB, Carpenter SR, Dahm CN, et al. (2001) Water in a changing world. Ecological Applications 11, 1027–1045.

Kalinowski ST, Meeuwig MH, Narum SR, Taper ML (2008) Stream trees: a statistical method for mapping genetic differences between populations of freshwater organisms to the sections of streams that connect them. Canadian Journal of Fisheries and Aquatic Sciences 65, 2752–2760.

Keyghobadi N (2007) The genetic implications of habitat fragmentation for animals. Canadian Journal of Zoology 85, 1049–1064.

Kingsford RT (2000) Ecological impacts of dams, water diversions and river management on floodplain wetlands in Australia. Austral Ecology 25, 109–127.

Lande R (1993) Risks of population extinction from demographic and environmental stochasticity and random catastrophes. American Naturalist, 911–927.

Landguth EL, Cushman SA, Schwartz MK, et al. (2010) Quantifying the lag time to detect barriers in landscape genetics. Molecular Ecology 19, 4179–4191.

Leblanc M, Tweed S, Van Dijk A, Timbal B (2012) A review of historic and future hydrological changes in the Murray-Darling Basin. Global and Planetary Change 80–81, 226–246.

Liermann CR, Nilsson C, Robertson J, Ng RY (2012) Implications of dam obstruction for global freshwater fish diversity. Bioscience 62, 539–548.

Luikart G, England PR, Tallmon D, Jordan S, Taberlet P (2003) The power and promise of population genomics: from genotyping to genome typing. Nature Reviews: Genetics 4, 981–994.

Manel S, Holderegger R (2013) Ten years of landscape genetics. Trends in Ecology & Evolution 28, 614–621.

Meirmans PG, Van Tienderen PH (2004) GenoType and GenoDive: two programs for the analysis of genetic diversity of asexual organisms. Molecular Ecology Notes 4, 792–794.

Morrissey MB, de Kerckhove DT (2009) The maintenance of genetic variation due to asymmetric gene flow in dendritic metapopulations. The American Naturalist 174, 875–889.

Murray–Darling Basin Authority (2013) Murray-Darling Basin Weir Information System. http://asdd.ga.gov.au/asdd/profileinfo/anzlic-jurisdic.xml#anzlic-jurisdic

Olden JD, Hogan ZS, Zanden MJV (2007) Small fish, big fish, red fish, blue fish: size-biased extinction risk of the world’s freshwater and marine fishes. Global Ecology and Biogeography 16, 694–701.

Pavlova A, Beheregaray LB, Coleman R, et al. (2017) Severe consequences of habitat fragmentation on genetic diversity of an endangered Australian freshwater fish: a call for assisted gene flow. Evolutionary Applications 10, 531–550.

Peterson B, Weber J, Kay E, Fisher H, Hoekstra H (2012) Double digest RADseq: an inexpensive method for de novo SNP discovery and genotyping in model and non-model species. PLoS ONE 7, e37135.

Puritz JB, Hollenbeck CM, Gold JR (2014) dDocent: a RADseq, variant-calling pipeline designed for population genomics of non-model organisms. PeerJ 2, e431.

Raj A, Stephens M, Pritchard JK (2014) fastSTRUCTURE: variational inference of population structure in large SNP data sets. Genetics 197, 573–589.

Ralls K, Ballou JD, Dudash MR, et al. (2018) Call for a paradigm shift in the genetic management of fragmented populations. Conservation Letters 11, e12412.

Saddlier S, Koehn JD, Hammer MP (2013) Let’s not forget the small fishes – conservation of two threatened species of pygmy perch in south-eastern Australia. Marine and Freshwater Research 64, 874–886.

Schielzeth H (2010) Simple means to improve the interpretability of regression coefficients. Methods in Ecology and Evolution 1, 103–113.

Stein J, Hutchinson M, Stein J (2014) A new stream and nested catchment framework for Australia. Hydrology and Earth System Sciences 18, 1917–1933.

Thomaz AT, Christie MR, Knowles LL (2016) The architecture of river networks can drive the evolutionary dynamics of aquatic populations. Evolution 70, 731–739.

Torterotot J-B, Perrier C, Bergeron NE, Bernatchez L (2014) Influence of forest road culverts and waterfalls on the fine-scale distribution of brook trout genetic diversity in a boreal watershed. Transactions of the American Fisheries Society 143, 1577–1591.

Wang IJ (2013) Examining the full effects of landscape heterogeneity on spatial genetic variation: a multiple matrix regression approach for quantifying geographic and ecological isolation. Evolution 67, 3403–3411.

Waples RS, Do C (2010) Linkage disequilibrium estimates of contemporary *N*e using highly variable genetic markers: a largely untapped resource for applied conservation and evolution. Evolutionary Applications 3, 244–262.

Webster MS, Colton MA, Darling ES, et al. (2017) Who should pick the winners of climate change? Trends in Ecology & Evolution.

Wedderburn S, Hammer M, Bice C (2012) Shifts in small-bodied fish assemblages resulting from drought-induced water level recession in terminating lakes of the Murray-Darling Basin, Australia. Hydrobiologia 691, 35–46.

Weeks AR, Stoklosa J, Hoffmann AA (2016) Conservation of genetic uniqueness of populations may increase extinction likelihood of endangered species: the case of Australian mammals. Frontiers in Zoology 13, 1.

Weir BS, Cockerham CC (1984) Estimating F-statistics for the analysis of population structure. Evolution 38, 1358–1370.

Weir BS, Hill W (2002) Estimating F-statistics. Annual Review of Genetics 36, 721–750.

Whiteley AR, Fitzpatrick SW, Funk WC, Tallmon DA (2015) Genetic rescue to the rescue. Trends in Ecology & Evolution 30, 42–49.

Wood JLA, Yates MC, Fraser DJ (2016) Are heritability and selection related to population size in nature? Meta-analysis and conservation implications. Evolutionary Applications 9, 640–657.

Yeomans KA, Golder PA (1982) The Guttman-Kaiser criterion as a predictor of the number of common factors. The Statistician 31, 221–229.

