## Supplementary Information for "Recent and rapid anthropogenic habitat fragmentation increases extinction risk for freshwater biodiversity"

Appendix S1. Library preparation and bioinformatics

Double digest RAD sequencing libraries were prepared following a protocol modified from Peterson et al. (2012). Using custom, in-house designed six-nucleotide barcodes with a minimum distance of three nucleotides 48 samples were multiplexed per Illumina lane. Samples were allocated to lanes randomly so that variation among sampling locations would not be confounded by artefacts arising due to differences among sequencing runs. For each lane of 48 individuals, 300ng total DNA from each sample was digested individually with two restriction enzymes *SbfI* and *MseI* (New England Biolabs) before ligation of the individual barcodes and RAD adapter sequences. Samples were then pooled into multiplex libraries of 12 uniquely barcoded individuals before purification with AMPure XP beads (Beckman Coulter Genomics) to remove any unligated adapters and adapter-to-adapter ligation products. Libraries were then size selected for an average of 500 base pair fragments (300-700bp) with a 1.5% Pippin prep electrophoresis gel (Sage Science) and quantified with a Qubit 2.0 fluorometer using the double-strand DNA broad range assay (Life Technologies). Polymerase chain reactions were then performed using two 25uL reactions per library (to reduce PCR bias associated with larger reaction volumes) before another bead purification step and fragment size evaluation with a Bioanalyzer 2100 (Agilent Technologies) using a DNA 7500 assay kit. Lastly, DNA quantity was assessed using real-time PCR to accurately equalize library concentrations before pooling four libraries of 12 samples together to create multiplex libraries of 48 uniquely barcoded samples for sequencing per one Illumina Hiseq2000 lane.

Libraries were sequenced as paired-end, 100-bp reads at the Genome Quebec/McGill University Innovation Center (Montreal, Canada). The raw data files were demultiplexed using the *process_radtags* component of *Stacks* v.1.04 (Catchen *et al.* 2011). Here, *process_radtags* was run twice utilizing the ‘rescue barcodes’ flag

(-r), initially to recover reads with up to two errors in the individual barcode sequences before subsequently using the same flag to recover reads with up to three errors in the RAD-tag. The demultiplexed sequences were then processed using *dDocent.FB* v.1.2 (Puritz *et al.* 2014). *dDocent* combines several existing bioinformatics software packages into a single pipeline and was designed specifically for paired-end RAD data and is thus able to take advantage of both forward and reverse reads for SNP discovery. The pipeline consists of four basic steps; quality trimming, de novo assembly of a catalogue of reference contigs, mapping of the trimmed reads to the reference catalogue, and variant calling. After demultiplexing, the raw reads were quality trimmed using *Trimmomatic* v.0.33 (Bolger *et al.* 2014). *Trimmomatic* was configured to remove adapter sequences and then remove low quality bases (PHRED <20) from the beginning and end of the reads, before using a sliding window approach (five nucleotide window) to trim the total read length if the average PHRED score drops below ten. Next *Rainbow* v.2.0.2 (Chong *et al.* 2012) was used to cluster reads based on similarity and then assemble the clusters into longer reference contigs with the maximum number of mismatches (-m) set to six. *CD-HIT* v.4.6 (Li *et al.* 2001) was then used to cluster the reference contigs based on 90% sequence similarity, retaining only the longest contig from each cluster in the final assembled reference contig data set. Using the MEM algorithm (Li 2013) implemented in *BWA* v.0.7.12-r1044 (Li & Durbin 2010), the quality trimmed reads were mapped to the reference contigs for each individual using default settings (match score -A=1, mismatch score -B=4, and gap-opening penalty -O=6). The resulting alignments were passed as indexed BAM files to the Bayesian variant caller *FreeBayes* v.0.9.20-8-gfef284a (Garrison & Marth 2012) to simultaneously detect SNPs, Indels, and more complex multi-nucleotide polymorphisms (MNPs) with default settings (minimum mapping quality -m=5, minimum base quality -q=5, and maximum complex gap -E=3). The resulting variant call file (VCF) containing information on sequence variation across all project samples was subsequently filtered using custom BASH scripts utilizing *VCFtools* (Danecek *et al.* 2011) and *vcflib*.

The following steps were implemented in order to filter SNPs likely to be the result of sequencing errors, paralogs, multi-copy loci and artefacts of library preparation and are based on scripts from the *dDocent* GitHub page (https://github.com/jpuritz/dDocent/). 1) Allele balance: for each locus, you should expect an approximately equal number of reads for the reference and alternate alleles for individuals called as heterozygotes. Loci were therefore removed if the proportion of alternate to reference allele was <0.25 or >0.75 across all heterozygote individuals. 2) Read orientation: each SNP should only occur in either forward or reverse reads. Those occurring in both are potentially paralogs and were accordingly filtered to retain only loci with at least 100 times more forward than reverse reads (or alternately, the opposite of 100 times more reverse than forward reads). 3) Mapping quality: as both alleles of a locus should start from the same RAD cut site, mapping quality scores (probability that the *BWA* alignment is correct) for the two alleles should be similar. Loci with a mapping quality score ratio (alternate allele mapping score/ reference allele mapping score) <90% or >110% were discarded. 4) Paired reads: loci where properly paired reads map to the reference allele but only unpaired reads map to the alternate allele are also indicative of potential paralogs and were removed. 5) Read quality: loci with overall low read quality scores (less than 25% of read depth) were discarded. Additionally, Li (2014), found a predictable relationship between Illumina read quality scores and read depth, such that where loci are covered by a high number of reads, quality scores are likely to be inflated. In this case, a higher quality score threshold is required to distinguish true variants from errors. Consequently, for loci with unusually high read depths (greater than the mean depth plus three times the square root of the mean), those with quality scores less than two times their read depth were also removed. 6) Read depth: finally, the read depth of each locus was recalculated and the frequency distribution of mean depth per locus, averaged over all individuals was assessed to identify and remove loci with abnormally high coverage.

### References


Bolger AM, Lohse M, Usadel B (2014) Trimmomatic: a flexible trimmer for Illumina sequence data. *Bioinformatics* **30**, 2114–2120.

Catchen JM, Amores A, Hohenlohe P, Cresko W, Postlethwait JH (2011) Stacks: building and genotyping loci de novo from short-read sequences. *G3: Genes, Genomes, Genetics* **1**, 171–182.

Chong Z, Ruan J, Wu C-I (2012) Rainbow: an integrated tool for efficient clustering and assembling RAD-seq reads. *Bioinformatics* **28**, 2732–2737.

Danecek P, Auton A, Abecasis G*, et al.* (2011) The variant call format and VCFtools. *Bioinformatics* **27**, 2156–2158.

Garrison E, Marth G (2012) Haplotype-based variant detection from short-read sequencing. *arXiv preprint arXiv:1207.3907*.

Li H (2013) Aligning sequence reads, clone sequences and assembly contigs with BWA-MEM. *arXiv preprint arXiv:1303.3997*.

Li H (2014) Toward better understanding of artifacts in variant calling from high-coverage samples. *Bioinformatics*.

Li H, Durbin R (2010) Fast and accurate long-read alignment with Burrows–Wheeler transform. *Bioinformatics* **26**, 589–595.

Li W, Jaroszewski L, Godzik A (2001) Clustering of highly homologous sequences to reduce the size of large protein databases. *Bioinformatics* **17**, 282–283.

Peterson B, Weber J, Kay E, Fisher H, Hoekstra H (2012) Double digest RADseq: an inexpensive method for de novo SNP discovery and genotyping in model and non-model species. *PLoS ONE* **7**, e37135.

Puritz JB, Hollenbeck CM, Gold JR (2014) dDocent: a RADseq, variant-calling pipeline designed for population genomics of non-model organisms. *PeerJ* **2**, e431.

Table S1. Pairwise population $F_{ST}$ among *Nannoperca australis* sampling sites ($F_{ST}$ below and *P* values above the diagonal).

| | TBA | ALE | MID | MUN | MCM | MIC | JHA | MER | TRA | YEA | PRA | SEV | BEN | SAM | LIM | KIN | HAP | MEA | GAP | ALB | SPR | GLE | TAL | COP | LRT |
|---|---|---|---|---|---|---|---|---|---|---|---|---|---|---|---|---|---|---|---|---|---|---|---|---|---|
| TBA | – | 0.001 | 0.001 | 0.001 | 0.001 | 0.001 | 0.001 | 0.001 | 0.001 | 0.002 | 0.001 | 0.001 | 0.001 | 0.001 | 0.001 | 0.001 | 0.001 | 0.001 | 0.001 | 0.001 | 0.001 | 0.001 | 0.002 | 0.001 | 0.001 |
| ALE | 0.035 | – | 0.001 | 0.003 | 0.001 | 0.001 | 0.001 | 0.001 | 0.001 | 0.001 | 0.001 | 0.001 | 0.001 | 0.001 | 0.001 | 0.001 | 0.001 | 0.001 | 0.001 | 0.001 | 0.001 | 0.001 | 0.001 | 0.001 | 0.001 |
| MID | 0.027 | 0.013 | – | 0.660 | 0.001 | 0.001 | 0.001 | 0.001 | 0.001 | 0.001 | 0.001 | 0.001 | 0.001 | 0.003 | 0.001 | 0.001 | 0.001 | 0.001 | 0.001 | 0.001 | 0.001 | 0.001 | 0.001 | 0.001 | 0.001 |
| MUN | 0.033 | 0.015 | -0.002 | – | 0.002 | 0.001 | 0.001 | 0.001 | 0.001 | 0.001 | 0.001 | 0.001 | 0.002 | 0.002 | 0.001 | 0.001 | 0.001 | 0.002 | 0.001 | 0.001 | 0.001 | 0.001 | 0.001 | 0.001 | 0.001 |
| MCM | 0.481 | 0.402 | 0.432 | 0.452 | – | 0.001 | 0.001 | 0.001 | 0.001 | 0.001 | 0.001 | 0.001 | 0.001 | 0.001 | 0.001 | 0.001 | 0.001 | 0.001 | 0.001 | 0.001 | 0.001 | 0.001 | 0.001 | 0.001 | 0.001 |
| MIC | 0.456 | 0.376 | 0.407 | 0.427 | 0.654 | – | 0.001 | 0.001 | 0.001 | 0.001 | 0.001 | 0.001 | 0.001 | 0.001 | 0.001 | 0.001 | 0.001 | 0.001 | 0.001 | 0.001 | 0.001 | 0.001 | 0.001 | 0.001 | 0.001 |
| JHA | 0.495 | 0.414 | 0.436 | 0.466 | 0.677 | 0.629 | – | 0.001 | 0.001 | 0.001 | 0.001 | 0.001 | 0.001 | 0.001 | 0.001 | 0.001 | 0.001 | 0.001 | 0.001 | 0.001 | 0.001 | 0.001 | 0.001 | 0.001 | 0.001 |
| MER | 0.623 | 0.547 | 0.585 | 0.609 | 0.747 | 0.712 | 0.725 | – | 0.001 | 0.001 | 0.001 | 0.001 | 0.001 | 0.001 | 0.001 | 0.001 | 0.001 | 0.001 | 0.001 | 0.001 | 0.001 | 0.001 | 0.001 | 0.001 | 0.001 |
| TRA | 0.582 | 0.501 | 0.539 | 0.564 | 0.74 | 0.699 | 0.719 | 0.171 | – | 0.001 | 0.001 | 0.001 | 0.001 | 0.001 | 0.001 | 0.001 | 0.001 | 0.001 | 0.001 | 0.001 | 0.001 | 0.001 | 0.001 | 0.001 | 0.001 |
| YEA | 0.541 | 0.458 | 0.493 | 0.517 | 0.719 | 0.674 | 0.698 | 0.128 | 0.111 | – | 0.001 | 0.001 | 0.001 | 0.001 | 0.001 | 0.001 | 0.001 | 0.002 | 0.001 | 0.001 | 0.001 | 0.001 | 0.001 | 0.001 | 0.001 |
| PRA | 0.216 | 0.163 | 0.167 | 0.17 | 0.438 | 0.392 | 0.395 | 0.399 | 0.349 | 0.299 | – | 0.001 | 0.001 | 0.001 | 0.001 | 0.001 | 0.001 | 0.001 | 0.001 | 0.001 | 0.001 | 0.001 | 0.001 | 0.001 | 0.001 |
| SEV | 0.237 | 0.176 | 0.178 | 0.184 | 0.453 | 0.422 | 0.437 | 0.559 | 0.517 | 0.479 | 0.152 | – | 0.001 | 0.001 | 0.001 | 0.001 | 0.001 | 0.001 | 0.001 | 0.001 | 0.001 | 0.001 | 0.001 | 0.001 | 0.001 |
| BEN | 0.217 | 0.158 | 0.158 | 0.162 | 0.428 | 0.388 | 0.406 | 0.538 | 0.493 | 0.457 | 0.13 | 0.115 | – | 0.002 | 0.001 | 0.001 | 0.001 | 0.001 | 0.001 | 0.001 | 0.001 | 0.001 | 0.001 | 0.001 | 0.001 |
| SAM | 0.206 | 0.145 | 0.148 | 0.151 | 0.429 | 0.388 | 0.399 | 0.532 | 0.487 | 0.449 | 0.118 | 0.107 | 0.035 | – | 0.001 | 0.001 | 0.001 | 0.001 | 0.001 | 0.001 | 0.001 | 0.001 | 0.001 | 0.001 | 0.001 |
| LIM | 0.471 | 0.399 | 0.421 | 0.437 | 0.623 | 0.59 | 0.594 | 0.687 | 0.672 | 0.653 | 0.382 | 0.352 | 0.212 | 0.253 | – | 0.001 | 0.001 | 0.001 | 0.001 | 0.001 | 0.001 | 0.001 | 0.001 | 0.001 | 0.001 |
| KIN | 0.513 | 0.421 | 0.448 | 0.479 | 0.683 | 0.662 | 0.696 | 0.759 | 0.751 | 0.733 | 0.506 | 0.501 | 0.487 | 0.479 | 0.661 | – | 0.001 | 0.001 | 0.001 | 0.001 | 0.001 | 0.001 | 0.001 | 0.001 | 0.001 |
| HAP | 0.455 | 0.353 | 0.38 | 0.407 | 0.668 | 0.637 | 0.68 | 0.756 | 0.746 | 0.721 | 0.442 | 0.436 | 0.42 | 0.419 | 0.636 | 0.332 | – | 0.001 | 0.001 | 0.001 | 0.001 | 0.001 | 0.002 | 0.001 | 0.001 |
| MEA | 0.378 | 0.287 | 0.301 | 0.328 | 0.599 | 0.579 | 0.623 | 0.715 | 0.693 | 0.662 | 0.38 | 0.375 | 0.361 | 0.353 | 0.592 | 0.153 | 0.173 | – | 0.001 | 0.001 | 0.001 | 0.001 | 0.001 | 0.001 | 0.001 |
| GAP | 0.435 | 0.361 | 0.38 | 0.389 | 0.594 | 0.586 | 0.62 | 0.701 | 0.677 | 0.651 | 0.427 | 0.425 | 0.416 | 0.408 | 0.601 | 0.599 | 0.566 | 0.502 | – | 0.001 | 0.001 | 0.001 | 0.001 | 0.001 | 0.001 |
| ALB | 0.339 | 0.278 | 0.287 | 0.288 | 0.509 | 0.501 | 0.53 | 0.629 | 0.593 | 0.564 | 0.339 | 0.339 | 0.331 | 0.321 | 0.528 | 0.527 | 0.48 | 0.42 | 0.089 | – | 0.001 | 0.001 | 0.001 | 0.001 | 0.001 |
| SPR | 0.488 | 0.417 | 0.436 | 0.447 | 0.65 | 0.639 | 0.669 | 0.74 | 0.72 | 0.696 | 0.475 | 0.479 | 0.467 | 0.46 | 0.648 | 0.656 | 0.63 | 0.57 | 0.39 | 0.202 | – | 0.001 | 0.001 | 0.001 | 0.001 |
| GLE | 0.505 | 0.433 | 0.453 | 0.466 | 0.664 | 0.649 | 0.679 | 0.746 | 0.729 | 0.706 | 0.489 | 0.492 | 0.48 | 0.472 | 0.656 | 0.668 | 0.642 | 0.585 | 0.409 | 0.22 | 0.051 | – | 0.001 | 0.001 | 0.001 |
| TAL | 0.462 | 0.39 | 0.409 | 0.418 | 0.644 | 0.625 | 0.659 | 0.74 | 0.717 | 0.69 | 0.451 | 0.458 | 0.441 | 0.434 | 0.639 | 0.646 | 0.613 | 0.549 | 0.373 | 0.185 | 0.101 | 0.096 | – | 0.001 | 0.001 |
| COP | 0.421 | 0.346 | 0.366 | 0.38 | 0.589 | 0.565 | 0.597 | 0.692 | 0.676 | 0.651 | 0.38 | 0.377 | 0.367 | 0.356 | 0.561 | 0.609 | 0.581 | 0.524 | 0.539 | 0.454 | 0.598 | 0.605 | 0.583 | – | 0.002 |
| LRT | 0.542 | 0.448 | 0.482 | 0.51 | 0.729 | 0.694 | 0.739 | 0.792 | 0.798 | 0.782 | 0.487 | 0.507 | 0.487 | 0.483 | 0.675 | 0.731 | 0.727 | 0.664 | 0.649 | 0.558 | 0.706 | 0.716 | 0.706 | 0.646 | – |

Table S2. Variation explained by retained environmental principal components and the correlation between the original variables and each component. Only variables significantly correlated (*P*<0.05) with each component are shown. Precipitation and disturbance variables shown in bold were analysed as individual variables.

| Category | PC | % Variation | Variables | Correlation |
|---|---|---|---|---|
| Temperature | Temp1 | 42.0% | STRCOLDMTHMIN | 0.895 |
| | | | STRWETQTEMP | 0.871 |
| | Temp2 | 33.3% | STRDRYQTEMP | 0.839 |
| | | | CATDRYQTEMP | 0.729 |
| Precipitation | – | | **CATDRYQRAIN** | – |
| | | | **STRWETQRAIN** | – |
| | | | **CATCOLDQRAIN** | – |
| Flow | Flow1 | 61.4% | CATEROSIVITY | 0.949 |
| | | | RUNPERENIALITY | 0.849 |
| | | | SUBEROSIVITY | 0.828 |
| | | | RUNANNMEAN | 0.569 |
| | | | RUNCVMAXMTH | -0.662 |
| | Flow2 | 21.7% | RUNANNMEAN | 0.762 |
| | | | RUNCVMAXMTH | 0.450 |
| | | | SUBEROSIVITY | -0.401 |
| Disturbance | – | | **FRDI** | – |
| | | | **CDI** | – |
| Topography | Topo1 | 52.1% | SUBELEMAX | 0.952 |
| | | | SUBELEMEAN | 0.879 |
| | | | CATELEMEAN | 0.729 |
| | | | CATSLOPE | 0.714 |
| | | | VALLEYSLOPE | 0.628 |
| | Topo2 | 31.2% | STRAHLER | 0.942 |
| | | | CATELEMEAN | 0.631 |
| | | | CATSLOPE | 0.447 |
| | | | VALLEYSLOPE | -0.568 |

Table S3. Simulated number of generations for global $F_{ST}$ to reach 0.2 for *Nannoperca australis* metapopulations of $N_e$=1000, $N_e$ =500 and $N_e$ =100 with increasing levels of habitat fragmentation. Simulations were based on a 1D stepping stone model assuming equal $N_e$ for each sub-population and were run for a burn-in of 20,000 generations with a migration rate of 0.5 between adjacent demes before 300 generations with no migration.

| Fragments | $N_e$ =1000 | $N_e$ =500 | $N_e$ =100 |
|---|---|---|---|
| 2 | 258 | 124 | 24 |
| 3 | 156 | 75 | 14 |
| 4 | 120 | 55 | 10 |
| 5 | 102 | 45 | 8 |
| 6 | 74 | 39 | 5 |
| 7 | 63 | 30 | 4 |
| 8 | 55 | 25 | 3 |
| 9 | 49 | 21 | 2 |
| 10 | 41 | 19 | 1 |

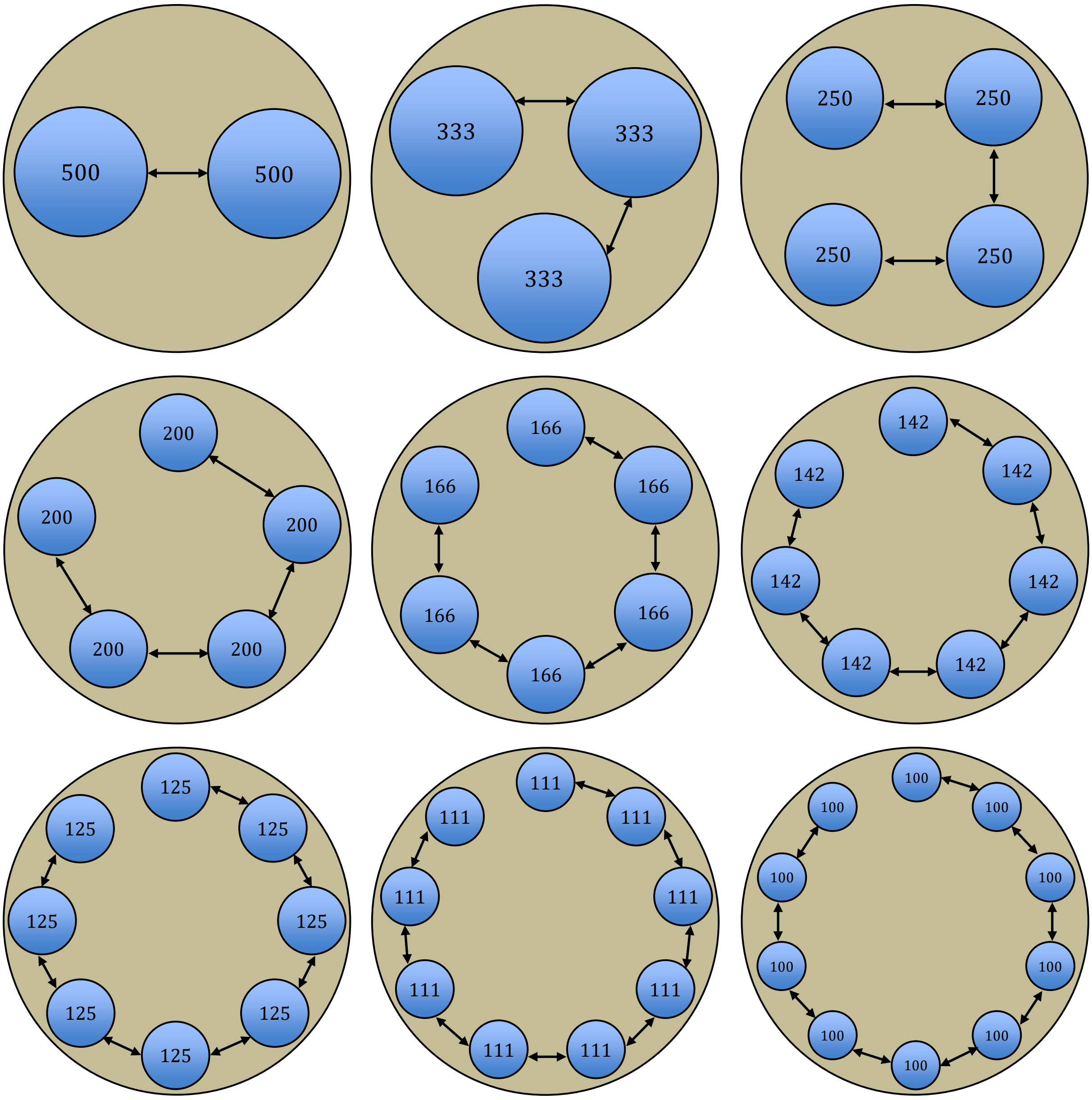


Figure S1. Schematic representation of simulated headwater catchment metapopulations for *Nannoperca australis*. Nine runs with increasing levels of fragmentation (n=2-10 habitat patches) were completed for each metapopulation. Each simulation was based on a stepping stone population model assuming equal $N_e$ for each sub-population, while maintaining a constant metapopulation $N_e$ of 1000 to simulate a concurrent reduction in habitat patch size with increasing fragmentation.

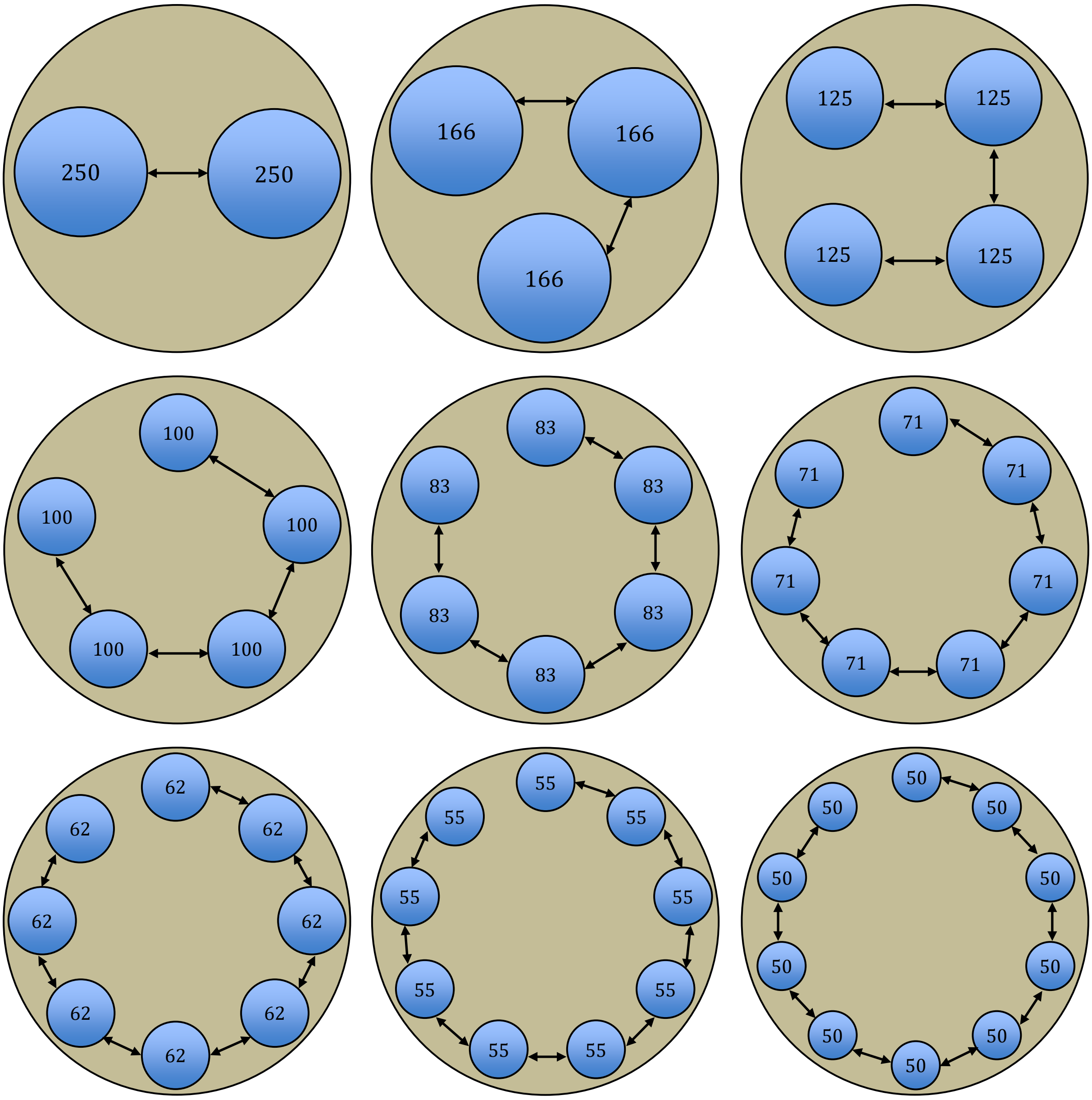


Figure S2. Schematic representation of simulated headwater catchment metapopulations for *Nannoperca australis*. Nine runs with increasing levels of fragmentation (n=2-10 habitat patches) were completed for each metapopulation. Each simulation was based on a stepping stone population model assuming equal $N_e$ for each sub-population, while maintaining a constant metapopulation $N_e$ of 500 to simulate a concurrent reduction in habitat patch size with increasing fragmentation.

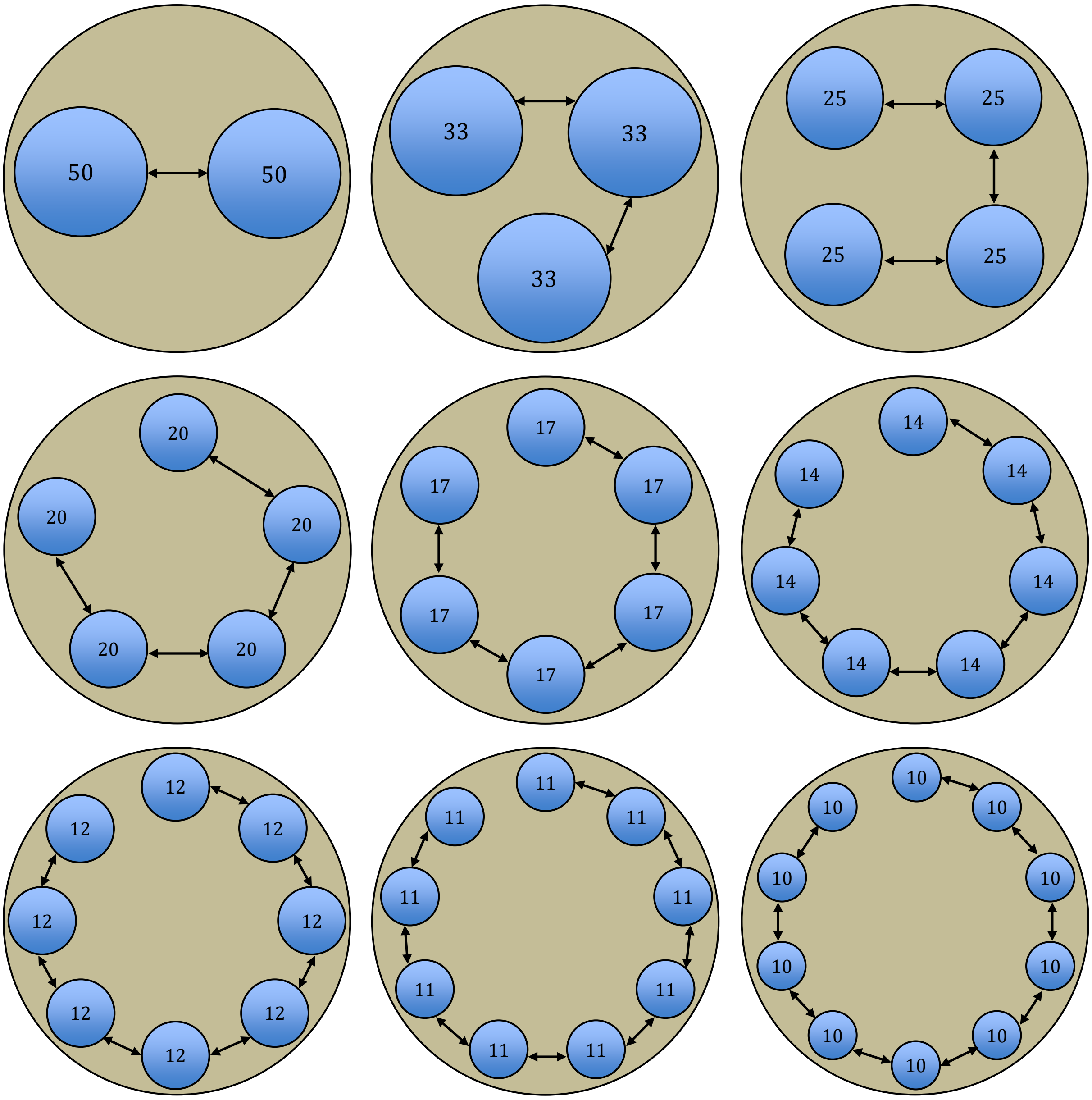


Figure S3. Schematic representation of simulated headwater catchment metapopulations for *Nannoperca australis*. Nine runs with increasing levels of fragmentation (n=2-10 habitat patches) were completed for each metapopulation. Each simulation was based on a stepping stone population model assuming equal $N_e$ for each sub-population, while maintaining a constant metapopulation $N_e$ of 100 to simulate a concurrent reduction in habitat patch size with increasing fragmentation.

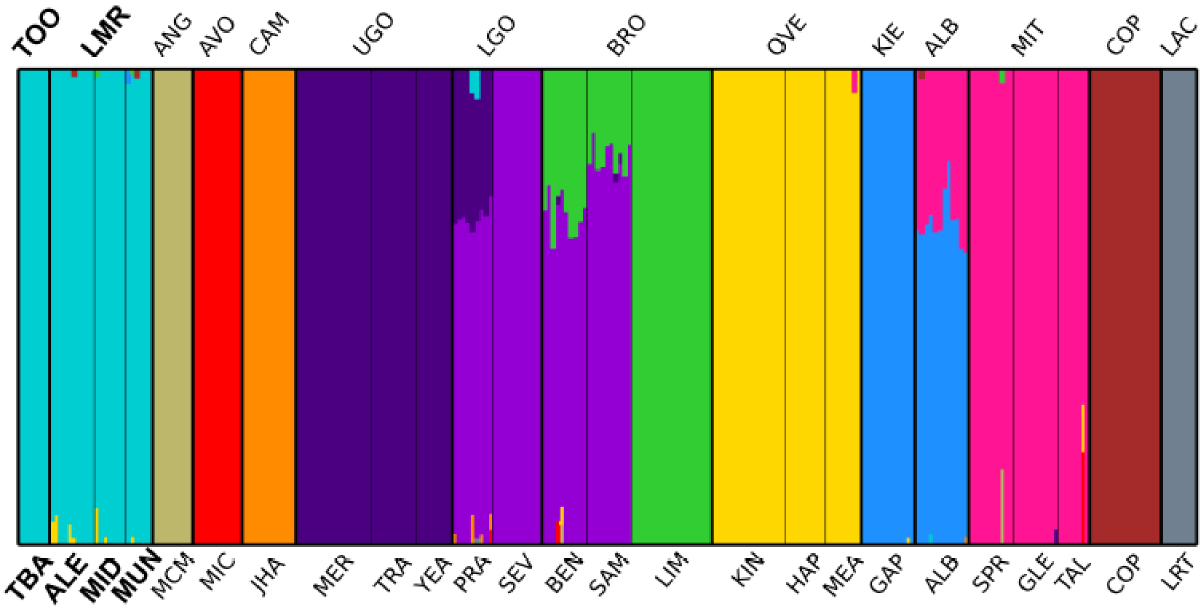


Figure S4. Admixture plot based on 3,443 SNPs for *Nannoperca australis* from the MDB depicting *K*=13 clusters determined by maximum marginal likelihood using *fastStructure*[29]. Codes above and below the plot refer to catchment and sampling site respectively (Table 1) Lowland wetland sites referred to as Lower Murray in the text are indicated in bold.

a)

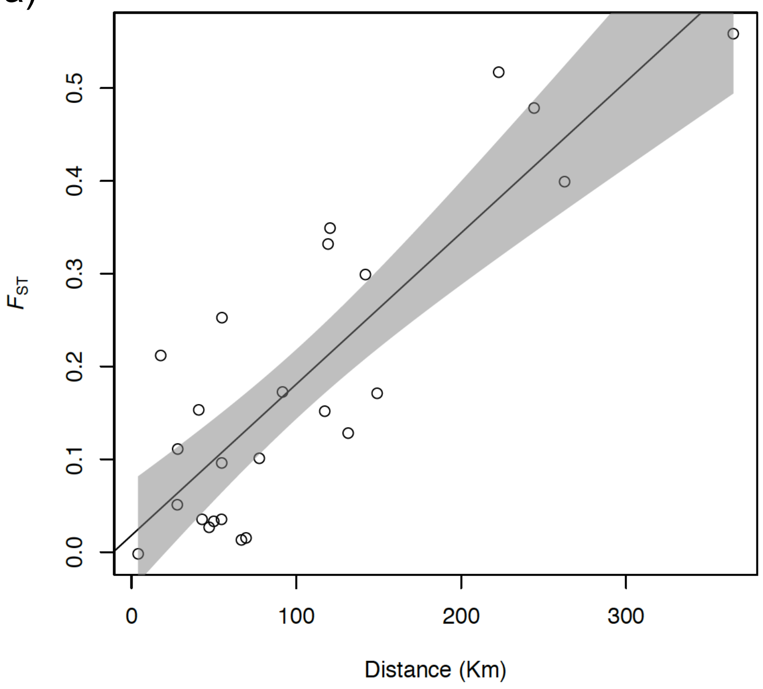


b)

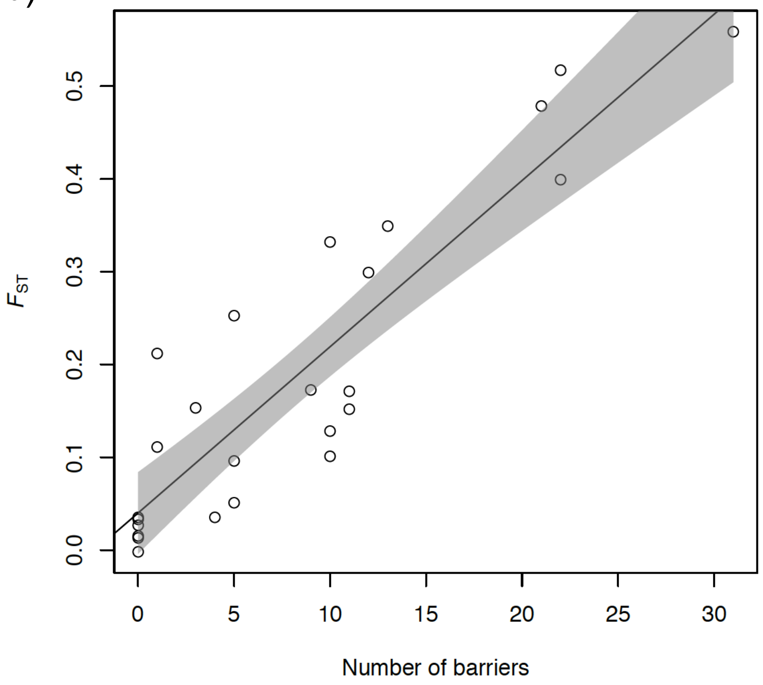


Figure S5. Plots of a) isolation by distance ($R^2$=0.73, β=0.002 [0.001- 0.002 95%CI], $P$<6.5X10$^{-08}$) and b) isolation by barrier ($R^2$=0.81, β=0.018 [0.014- 0.022 95%CI], $P$<6.5X10$^{-08}$) for *Nannoperca australis* in the Murray-Darling Basin considering only pairwise comparisons within catchments. Shaded areas represent the 95% confidence intervals.

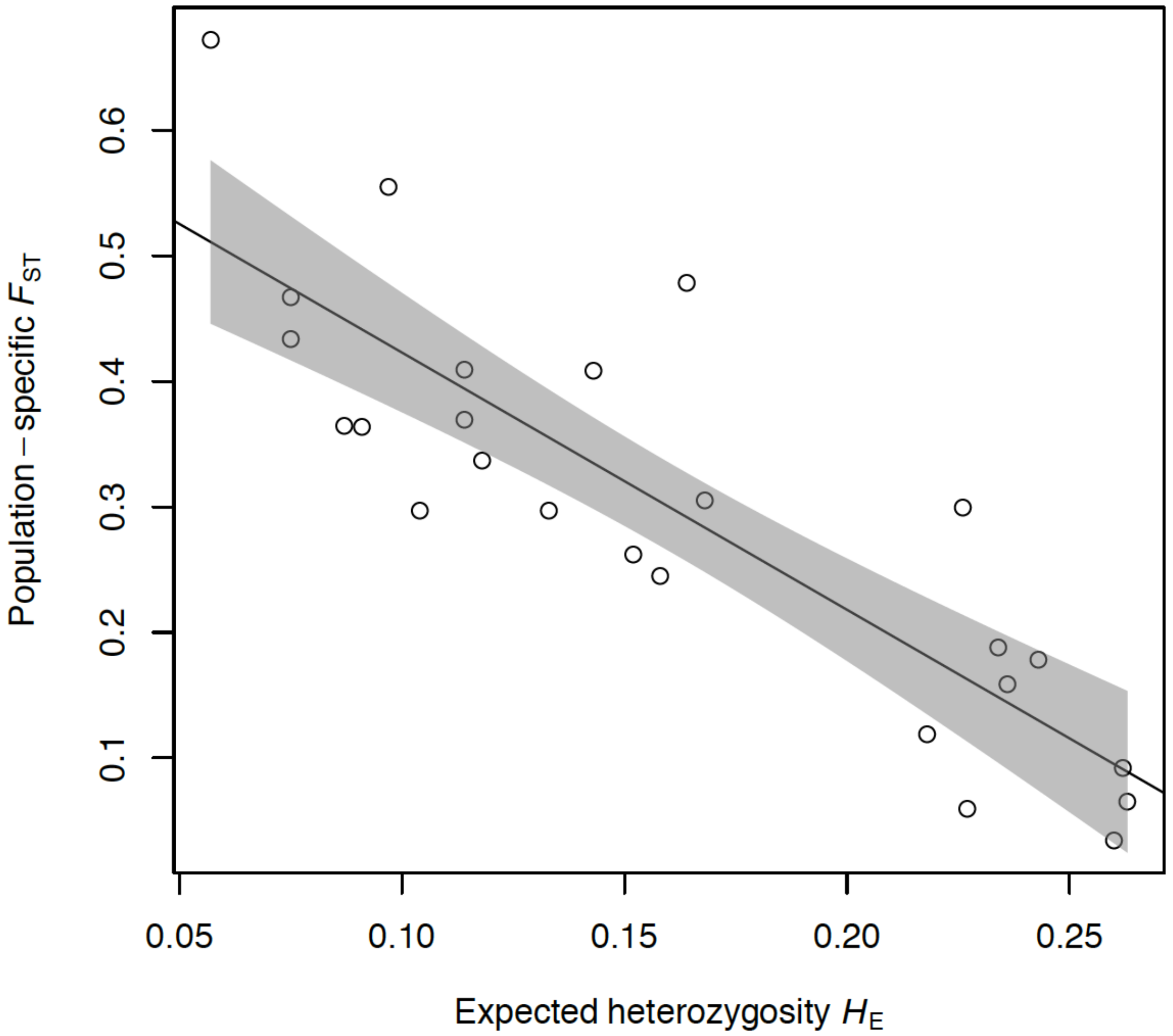


Figure S6. Regression plot of population-specific $F_{ST}$ vs. expected heterozygosity ($H_E$) ($R^2$=0.737, β=-2.05 [-2.58- -1.52 95%CI], $P$<1x10$^{-7}$) for populations of *Nannoperca australis*. Shaded area represents the 95% confidence interval.

Appendix S2. Metapopulation simulations

Simulations were run for three metapopulation sizes ($N_e$=1000, $N_e$=500 and $N_e$=100) using *SLiM 3.1* (Haller & Messer, 2018). Each simulation was based on a 1D stepping stone population model assuming equal $N_e$ for each sub-population and maintaining a constant metapopulation size to simulate a concurrent reduction in habitat patch size with increasing fragmentation. Each simulation consisted of four 100Kb genomic elements and assumed a constant mutation rate of $10^{-7}$ and recombination rate of $10^{-8}$. The simulations were run for 20,000 generations with a migration rate of 0.5 between adjacent sub-populations to reach migration–drift equilibrium, before simulating the construction of barriers by setting the migration rate to zero for 300 generations. Nine models with an increasing number of demes (2-10) were simulated for each metapopulation to examine the effect of increasing the number of barriers (Supplementary Fig. 3-5), and 100 replicate runs of each scenario were completed. The --weir-fst-pop command of VCFtools (Danecek et al., 2011) was used to calculate $F_{ST}$ for each replicate. To estimate the time required to reach current levels of population differentiation (assuming a generation time of one year [Humphries, 1995]) for each scenario, the number of generations (mean of the 100 replicates) needed to achieve $F_{ST}$=0.2 (mean contemporary $F_{ST}$ within upper Murray catchments is 0.196; Table S1) was plotted against the number of fragments for each simulation for the three metapopulation models. Scripts used to perform the simulations and analyses are available on Dryad: TBA. Although our assumption of $F_{ST}$=~0 before anthropogenic fragmentation almost certainly underestimates natural historical population structure, this figure provides the most conservative approach by maximizing the number of generations required to evolve current levels of differentiation.

**Appendix S3**. Plots of $F_{ST}$ vs. number of generations for SLiM 3.1 simulated metapopulations where $N_e$=1000. Scenarios were simulated with 2-10 sub-populations and each scenario was replicated 100 times. For each plot, the blue line represents a second order polynomial model of mean $F_{ST}$ calculated every ten generations, and the red star indicates the generation number at which $F_{ST}$ reaches 0.2 as calculated by applying the *predict R* function to the model. Error bars depict 95% confidence intervals.

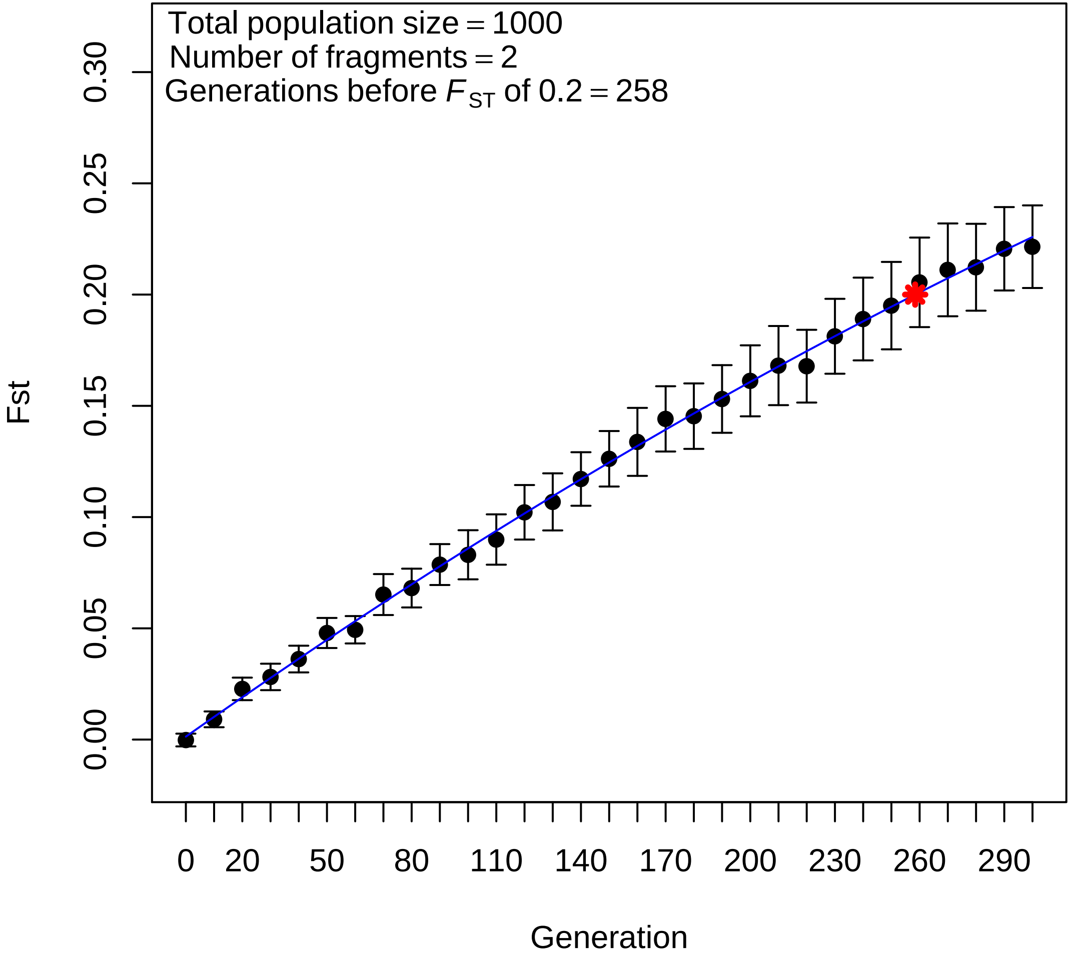

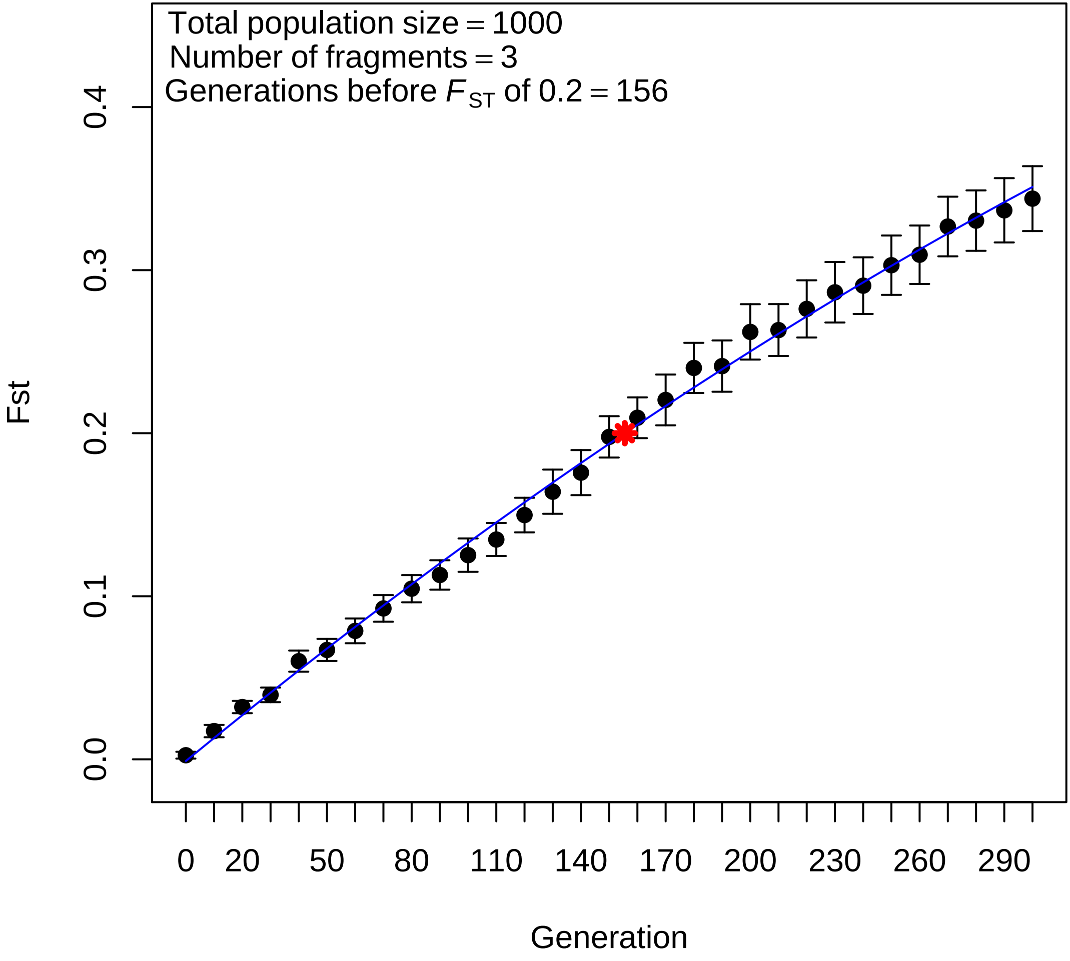

Total population size = 1000
Number of fragments = 3
Generations before $F_{ST}$ of 0.2 = 156
Fst
0.0
0.1
0.2
0.3
0.4
0
20
50
80
110
140
170
200
230
260
290
Generation

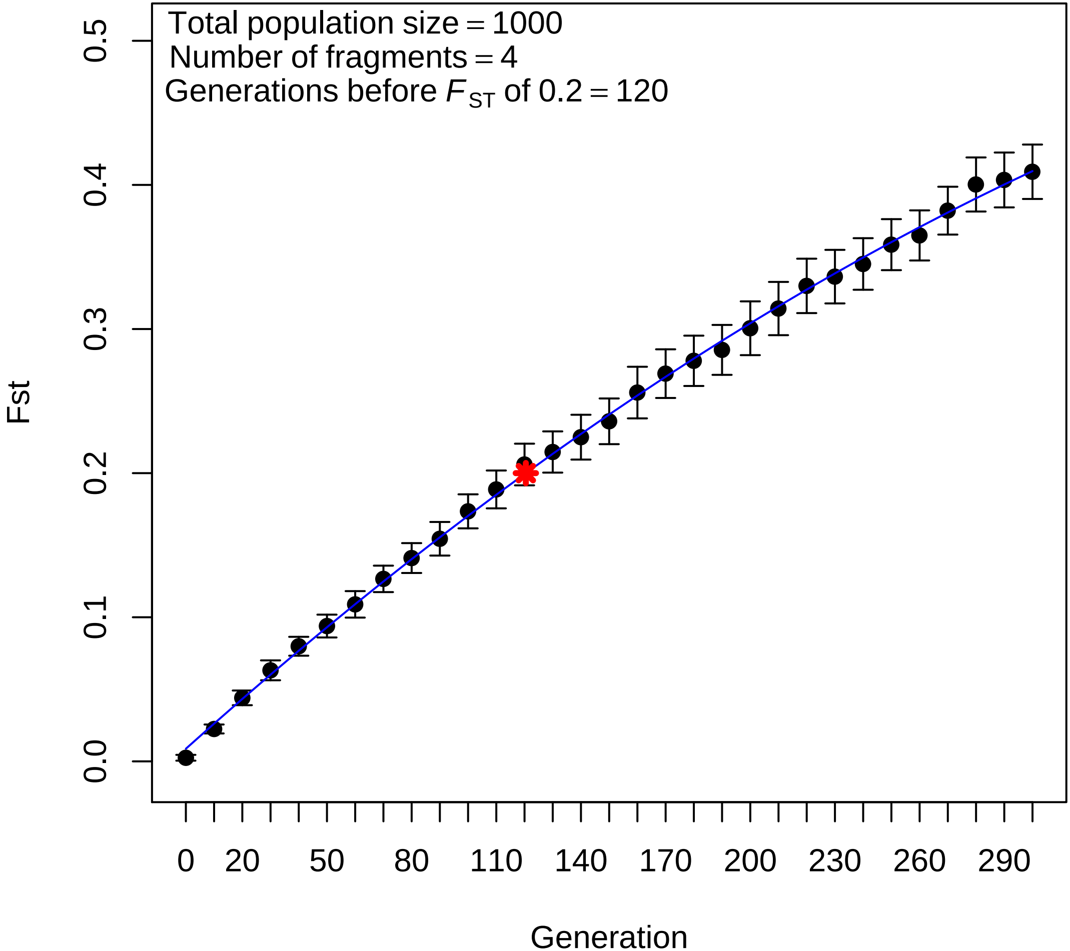

Total population size = 1000
Number of fragments = 4
Generations before $F_{ST}$ of 0.2 = 120
Fst
0.0
0.1
0.2
0.3
0.4
0.5
0
20
50
80
110
140
170
200
230
260
290
Generation

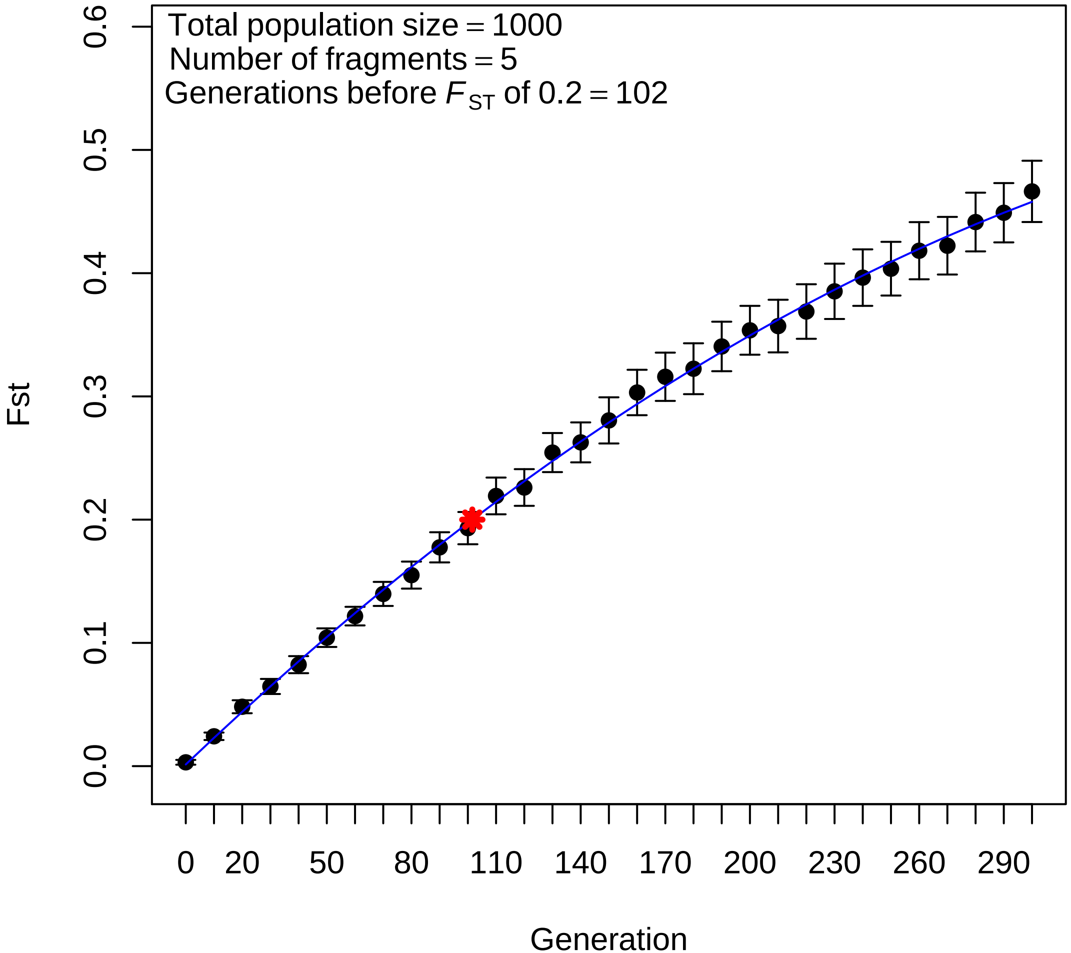

Total population size = 1000
Number of fragments = 5
Generations before $F_{ST}$ of 0.2 = 102
Fst
0.0
0.1
0.2
0.3
0.4
0.5
0.6
0
20
50
80
110
140
170
200
230
260
290
Generation

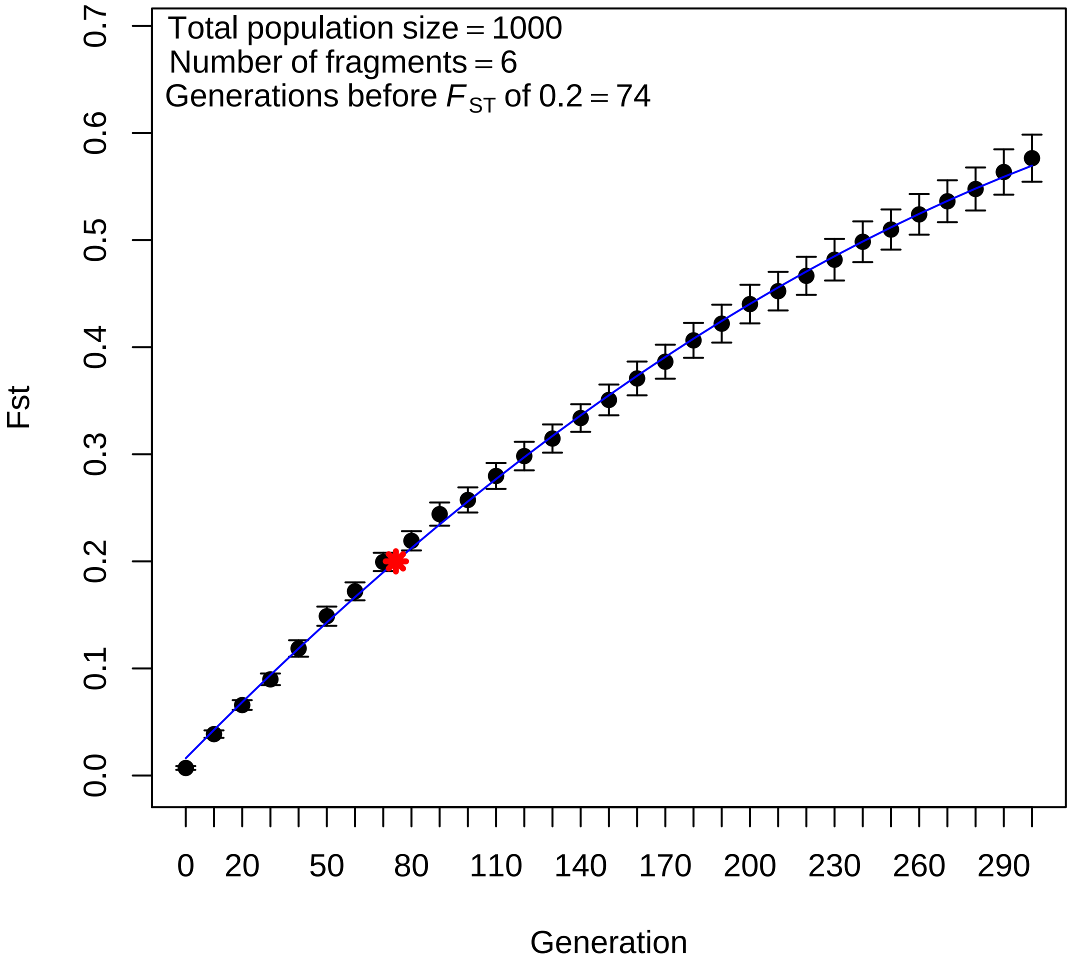
Total population size = 1000
Number of fragments = 6
Generations before $F_{ST}$ of 0.2 = 74
Fst
0.0
0.1
0.2
0.3
0.4
0.5
0.6
0.7
0
20
50
80
110
140
170
200
230
260
290
Generation

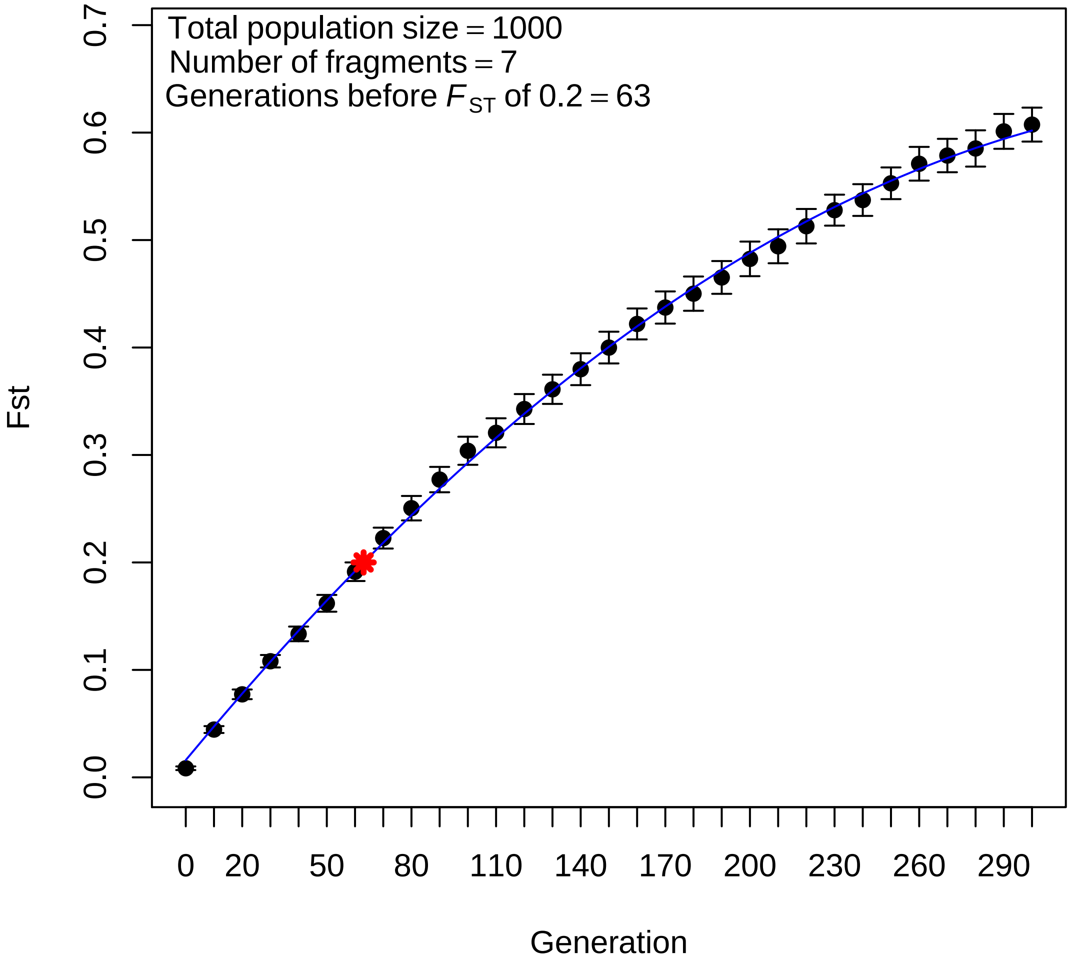
Total population size = 1000
Number of fragments = 7
Generations before $F_{ST}$ of 0.2 = 63
Fst
Generation
0.0
0.1
0.2
0.3
0.4
0.5
0.6
0.7
0
20
50
80
110
140
170
200
230
260
290

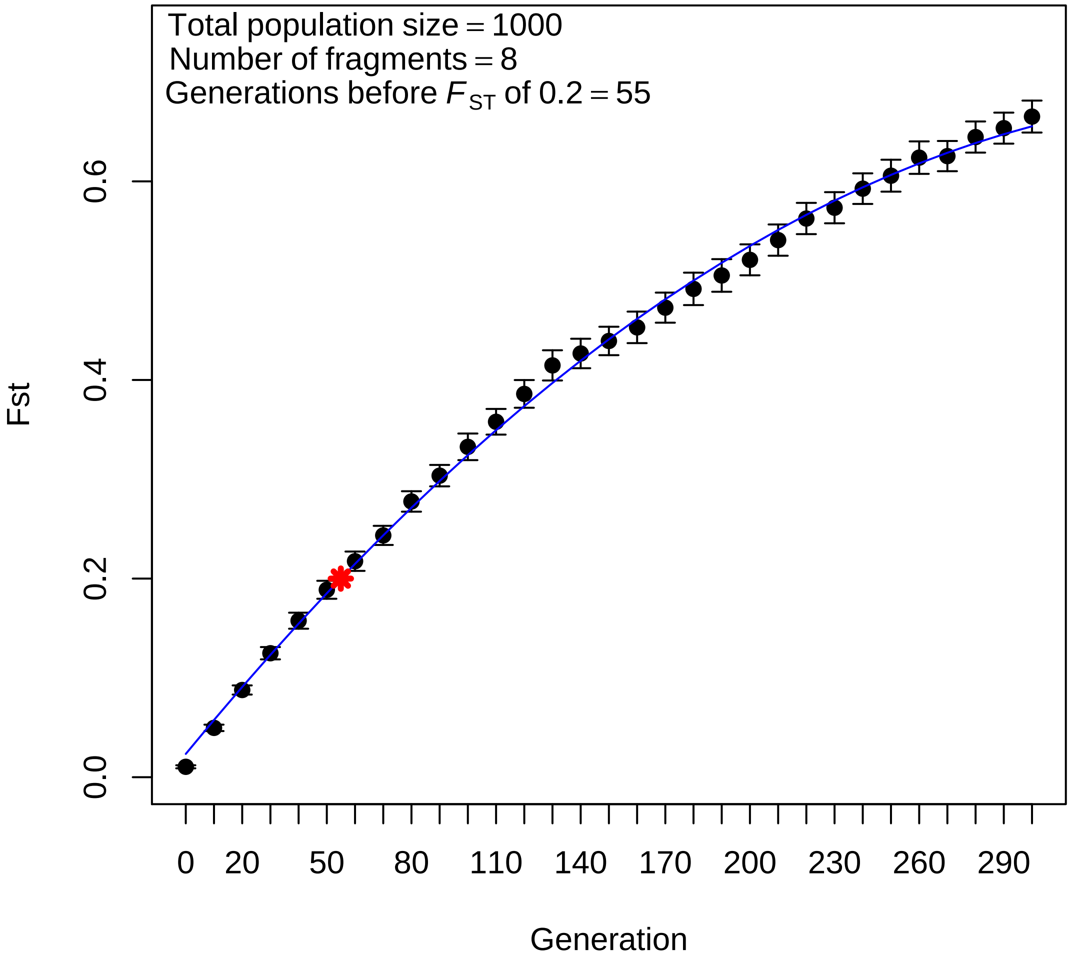

Total population size = 1000
Number of fragments = 8
Generations before $F_{ST}$ of 0.2 = 55
Fst
0.0
0.2
0.4
0.6
0
20
50
80
110
140
170
200
230
260
290
Generation

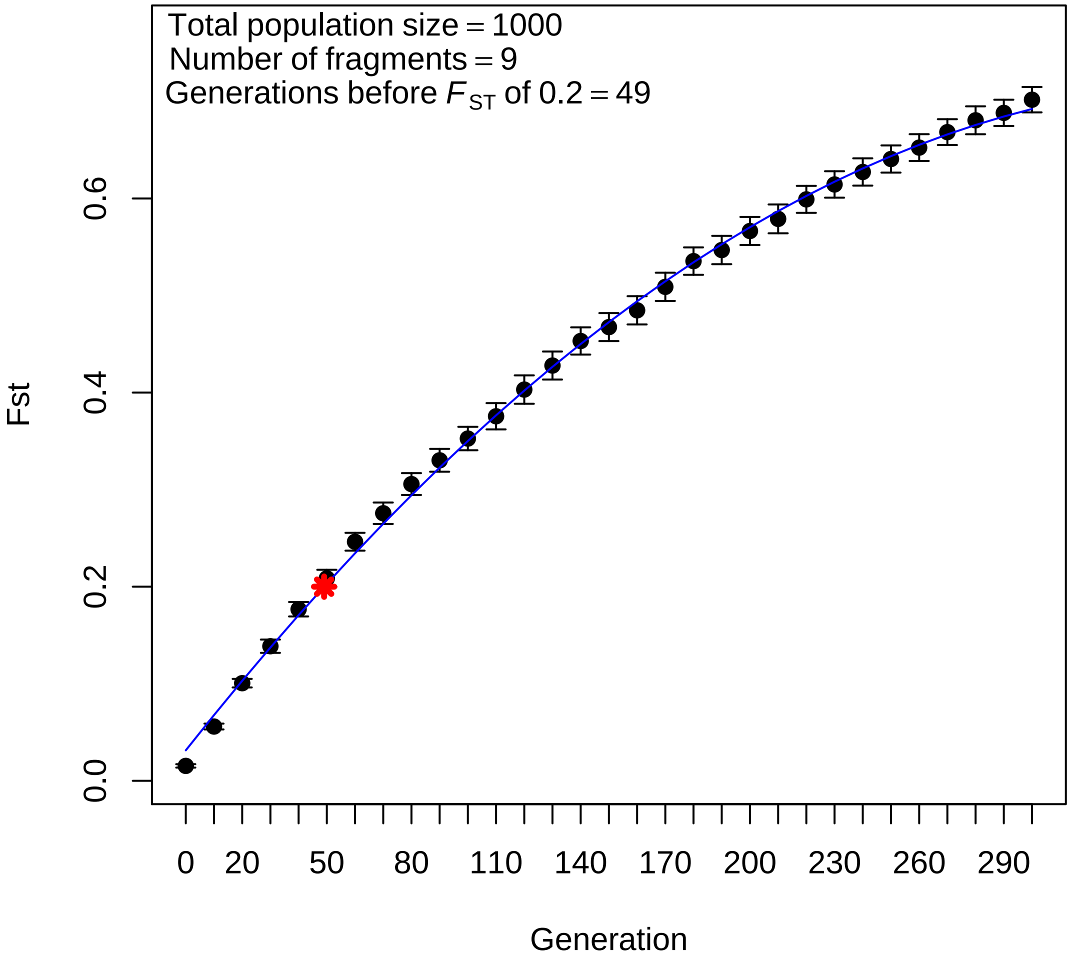

Total population size = 1000
Number of fragments = 9
Generations before $F_{ST}$ of 0.2 = 49
Fst
0.0
0.2
0.4
0.6
0
20
50
80
110
140
170
200
230
260
290
Generation

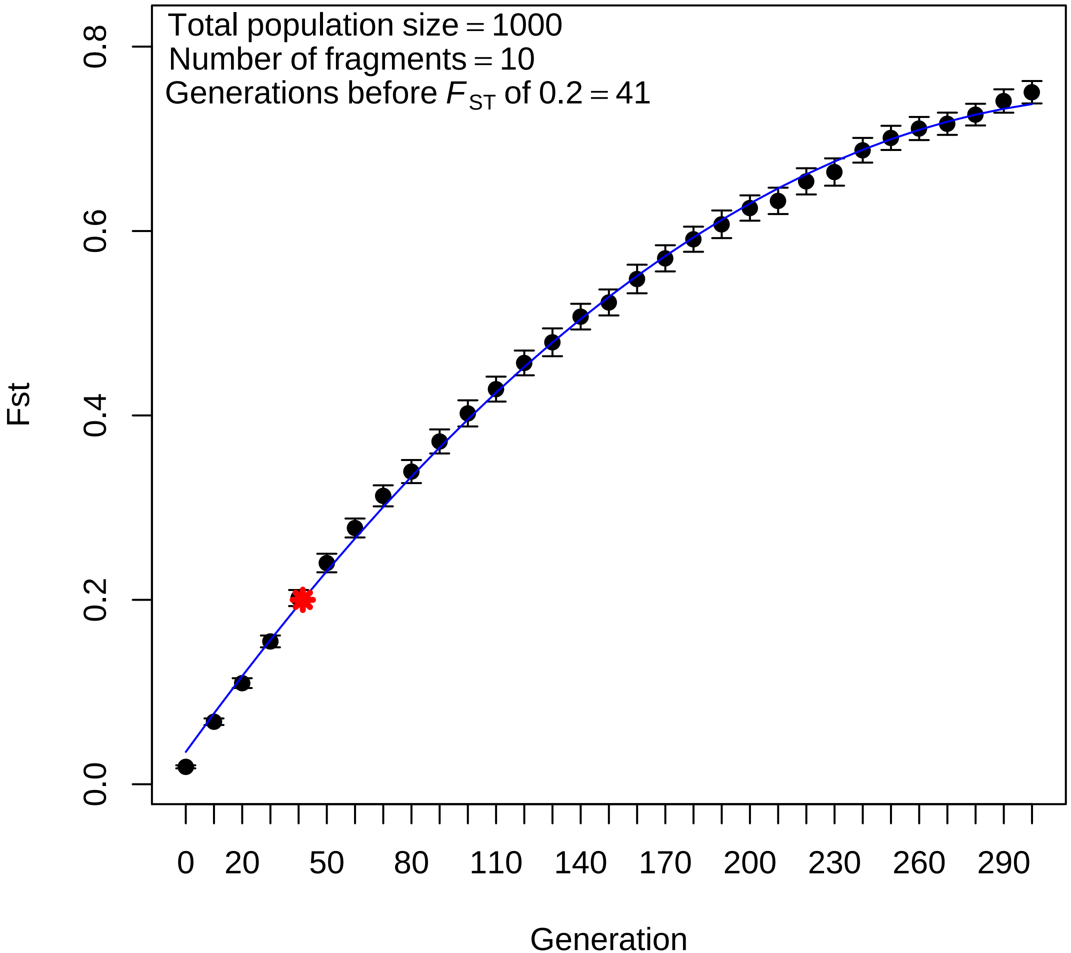

Total population size = 1000
Number of fragments = 10
Generations before $F_{ST}$ of 0.2 = 41
Fst
0.0
0.2
0.4
0.6
0.8
0
20
50
80
110
140
170
200
230
260
290
Generation

**Appendix S4**. Plots of $F_{ST}$ vs. number of generations for SLiM 3.1 simulated metapopulations where $N_e$=500. Scenarios were simulated with 2-10 sub-populations and each scenario was replicated 100 times. For each plot, the blue line represents a second order polynomial model of mean $F_{ST}$ calculated every ten generations, and the red star indicates the generation number at which $F_{ST}$ reaches 0.2 as calculated by applying the *predict R* function to the model. Error bars depict 95% confidence intervals.

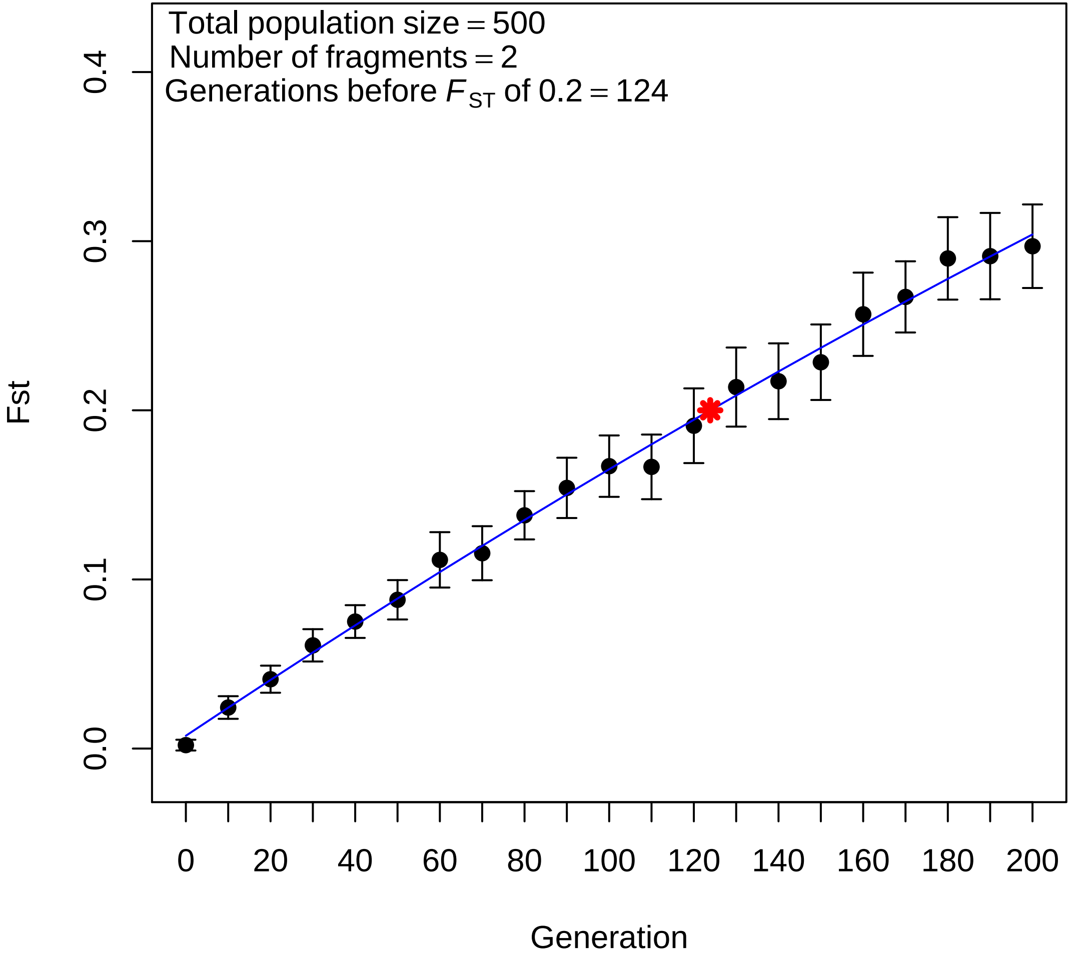

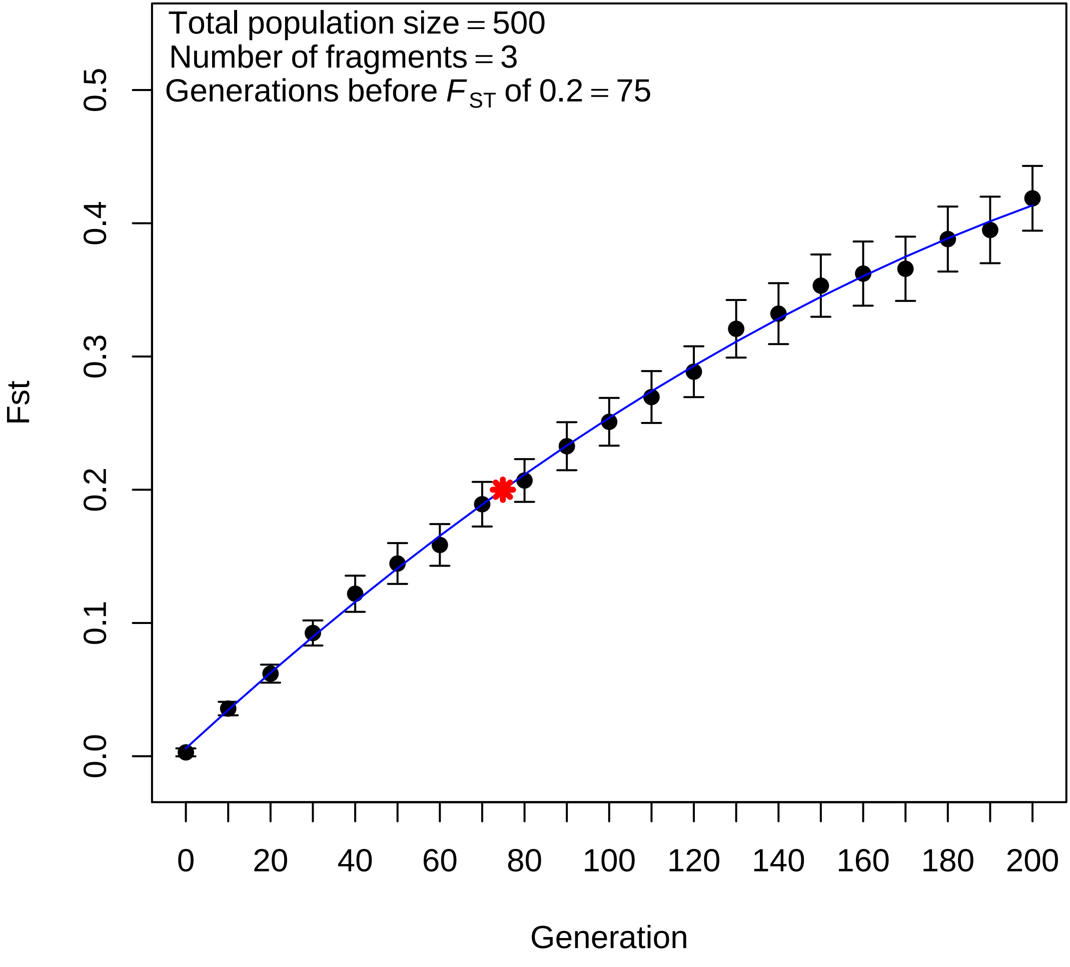

Total population size = 500
Number of fragments = 3
Generations before $F_{ST}$ of 0.2 = 75
Fst
0.0
0.1
0.2
0.3
0.4
0.5
0
20
40
60
80
100
120
140
160
180
200
Generation

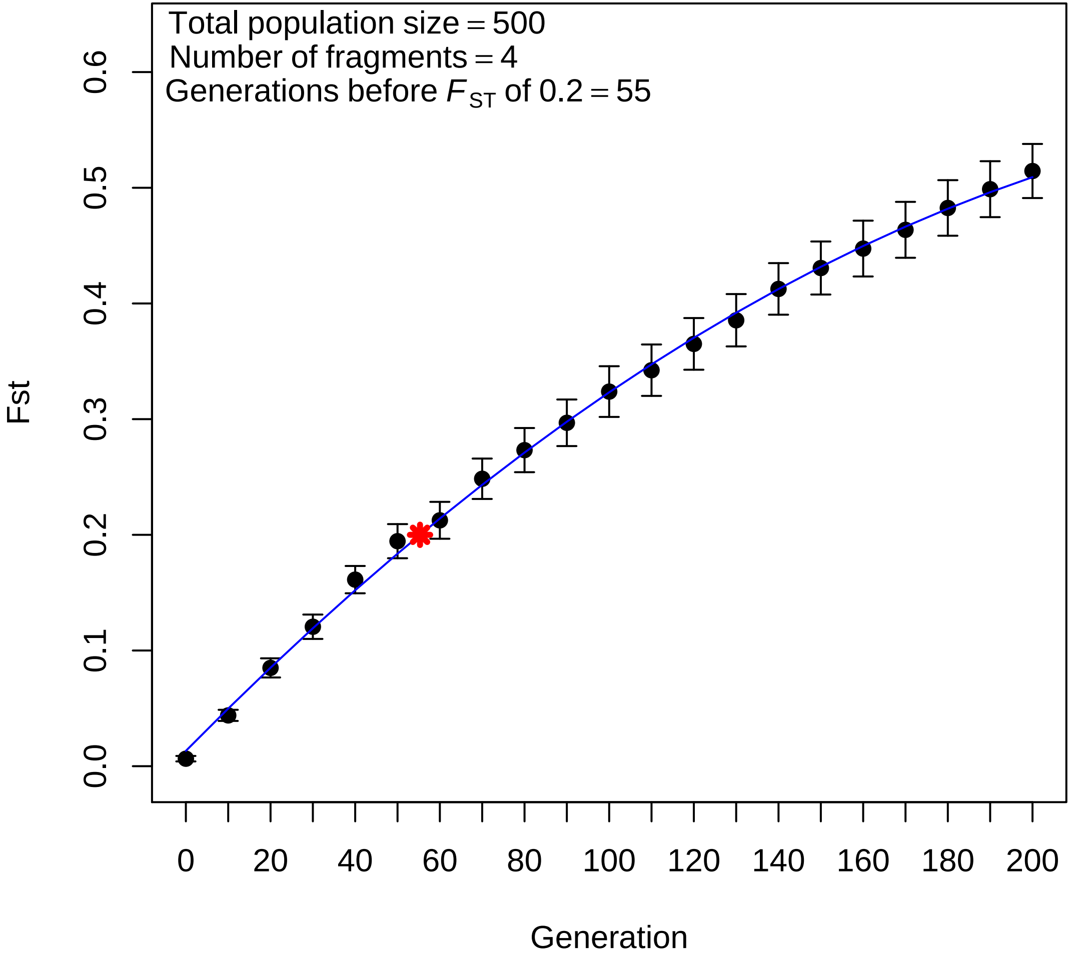

Total population size = 500
Number of fragments = 4
Generations before $F_{ST}$ of 0.2 = 55
Fst
Generation
0.0
0.1
0.2
0.3
0.4
0.5
0.6
0
20
40
60
80
100
120
140
160
180
200

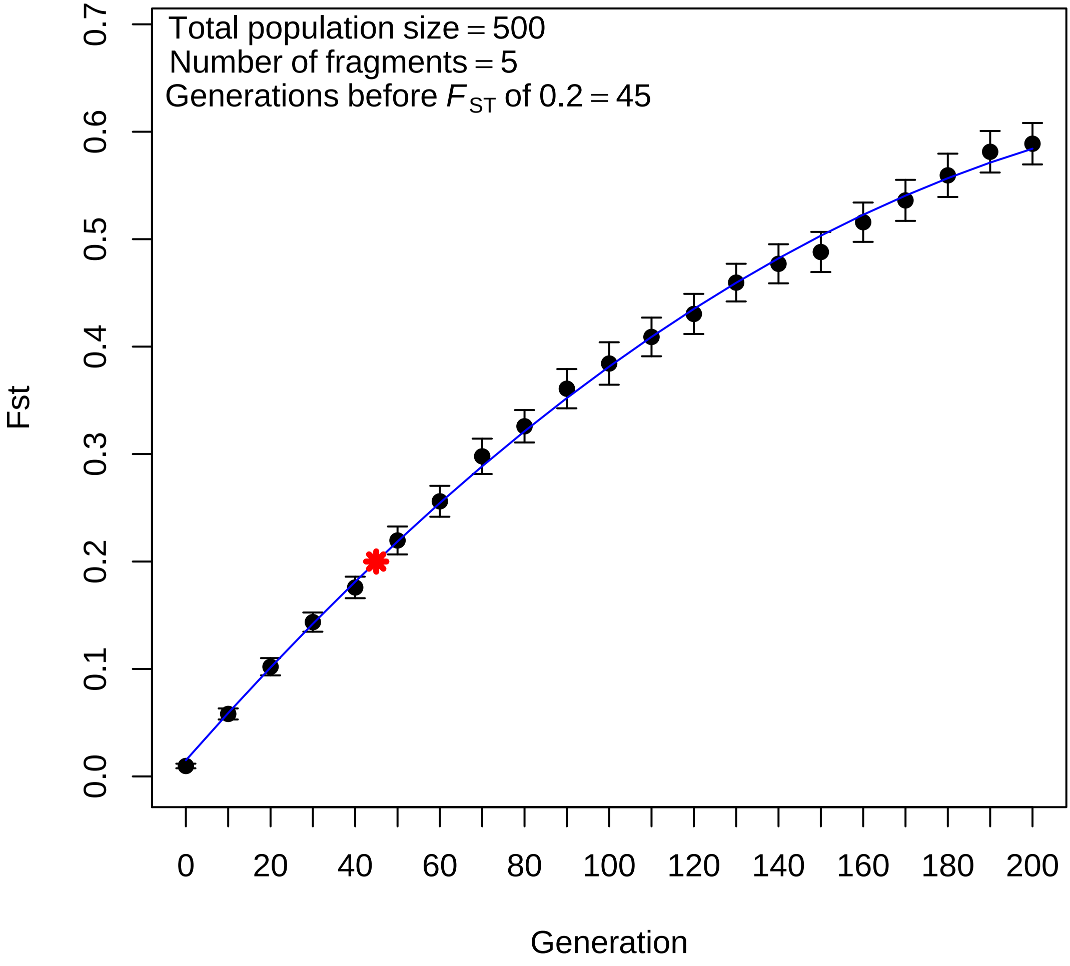

Total population size = 500
Number of fragments = 5
Generations before $F_{ST}$ of 0.2 = 45
Fst
0.0
0.1
0.2
0.3
0.4
0.5
0.6
0.7
0
20
40
60
80
100
120
140
160
180
200
Generation

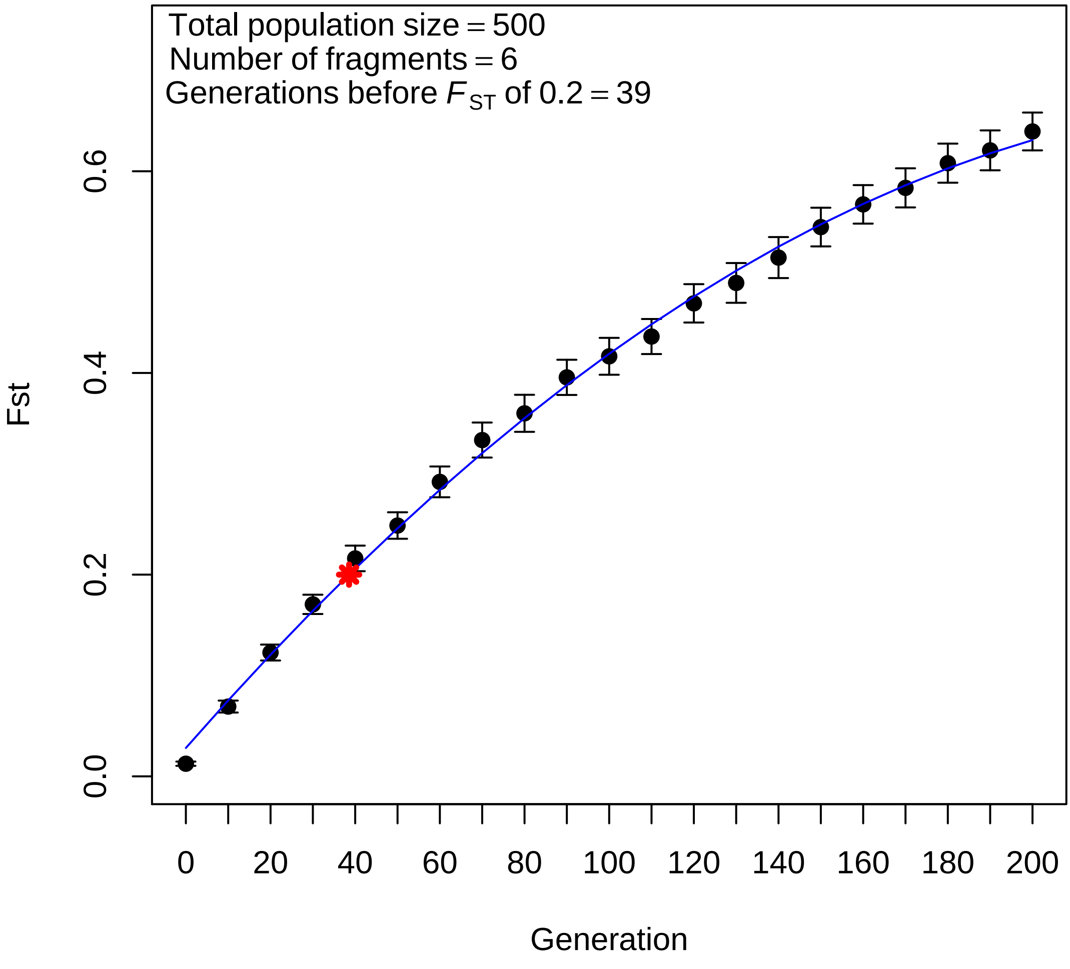

Total population size = 500
Number of fragments = 6
Generations before $F_{ST}$ of 0.2 = 39
Fst
0.0
0.2
0.4
0.6
0
20
40
60
80
100
120
140
160
180
200
Generation

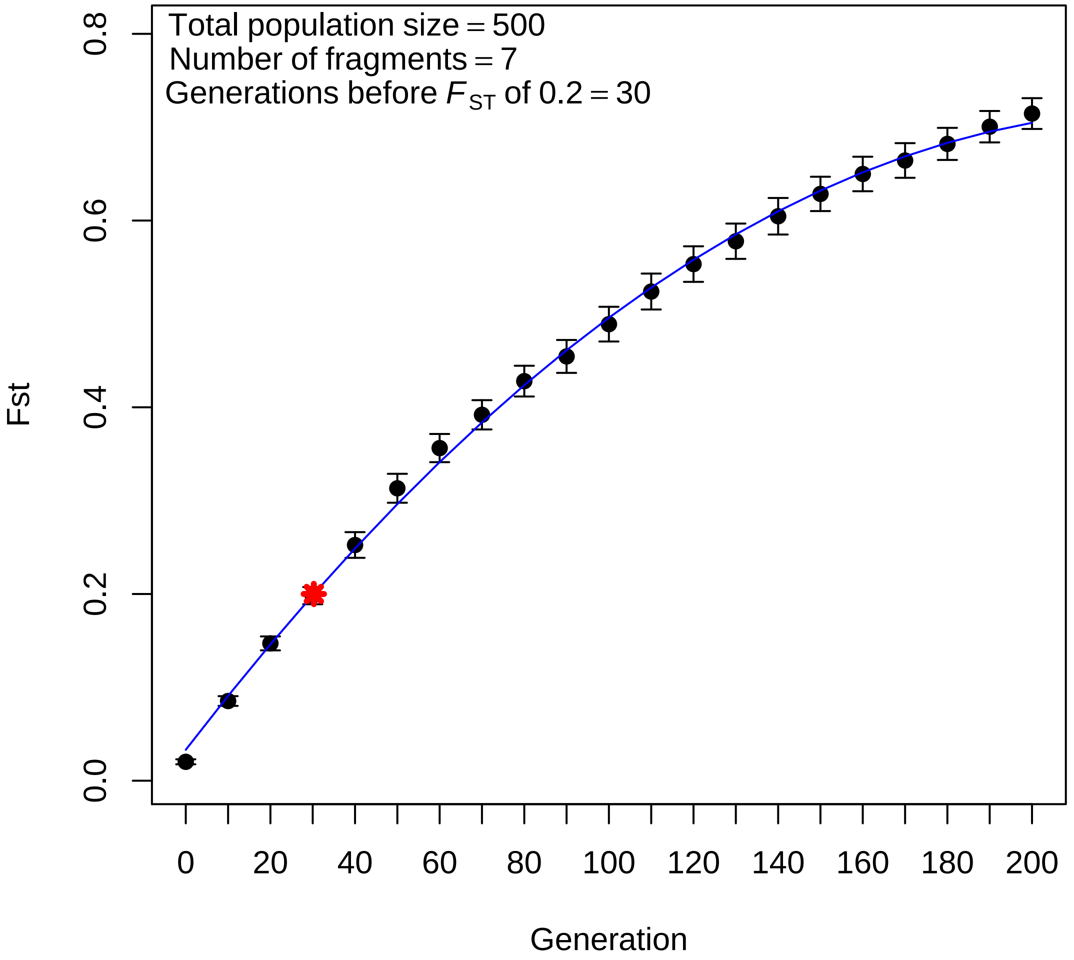

Total population size = 500
Number of fragments = 7
Generations before $F_{ST}$ of 0.2 = 30
Fst
0.0
0.2
0.4
0.6
0.8
0
20
40
60
80
100
120
140
160
180
200
Generation

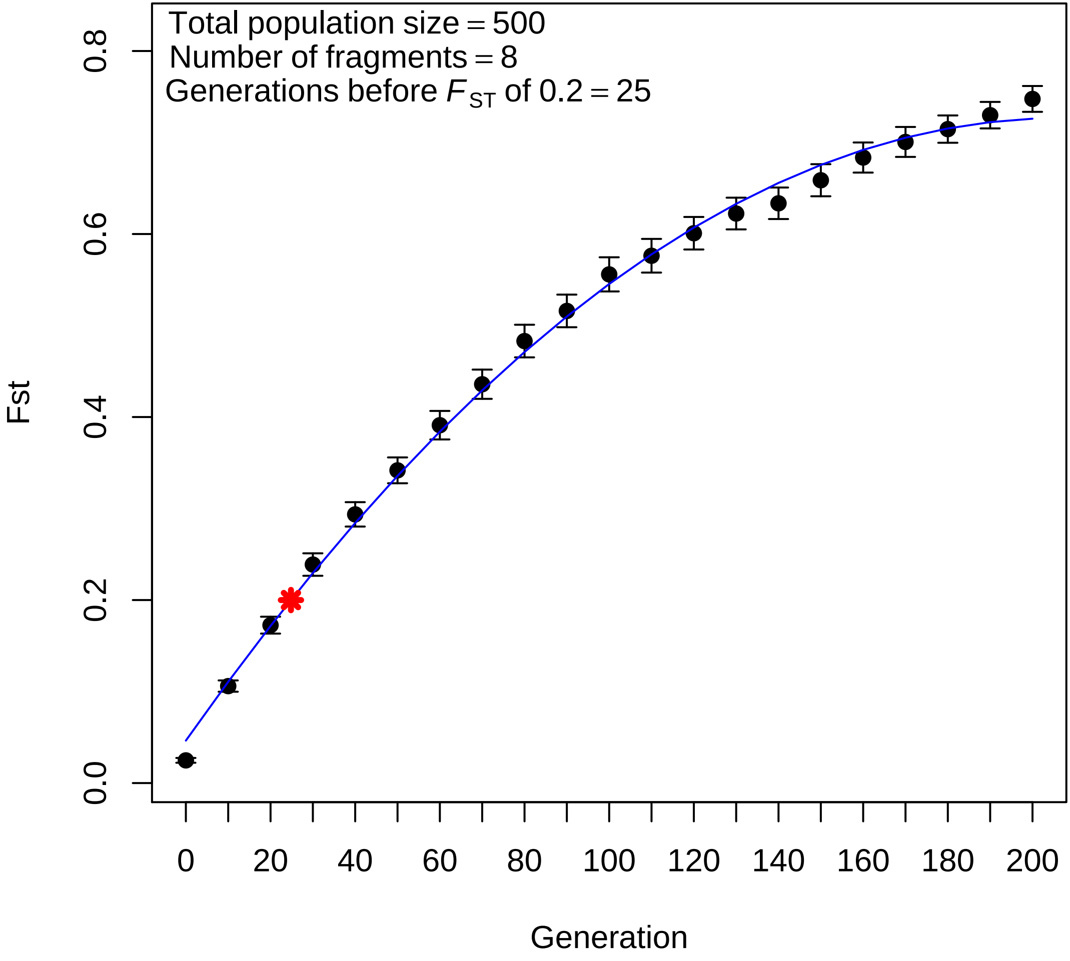

Total population size = 500
Number of fragments = 8
Generations before $F_{ST}$ of 0.2 = 25
Fst
0.0
0.2
0.4
0.6
0.8
0
20
40
60
80
100
120
140
160
180
200
Generation

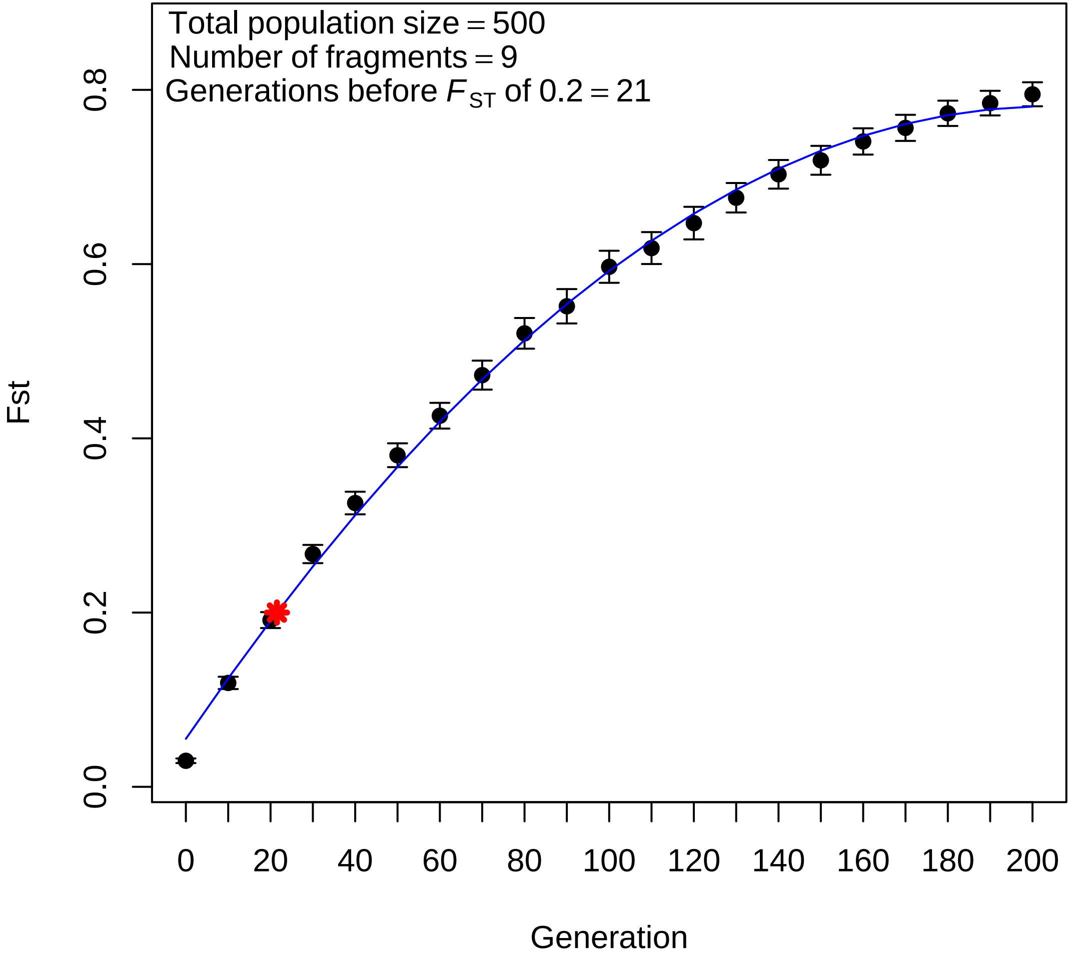

Total population size = 500
Number of fragments = 9
Generations before $F_{ST}$ of 0.2 = 21
Fst
0.0
0.2
0.4
0.6
0.8
0
20
40
60
80
100
120
140
160
180
200
Generation

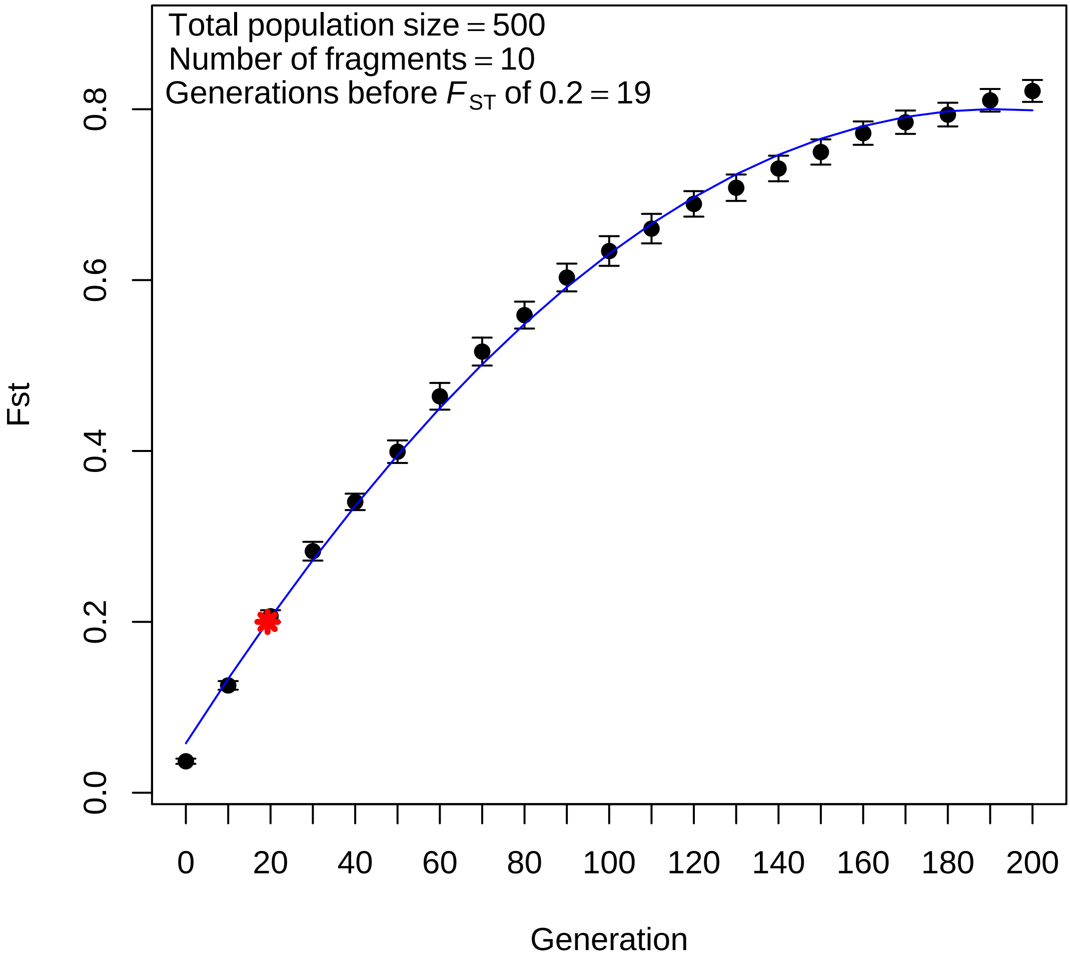

Total population size = 500
Number of fragments = 10
Generations before $F_{ST}$ of 0.2 = 19
Fst
0.0
0.2
0.4
0.6
0.8
0
20
40
60
80
100
120
140
160
180
200
Generation

**Appendix S5**. Plots of $F_{ST}$ vs. number of generations for SLiM 3.1 simulated metapopulations where $N_e$=100. Scenarios were simulated with 2-10 sub-populations and each scenario was replicated 100 times. For each plot, the blue line represents a second order polynomial model of mean $F_{ST}$ calculated every ten generations, and the red star indicates the generation number at which $F_{ST}$ reaches 0.2 as calculated by applying the *predict R* function to the model. Error bars depict 95% confidence intervals.

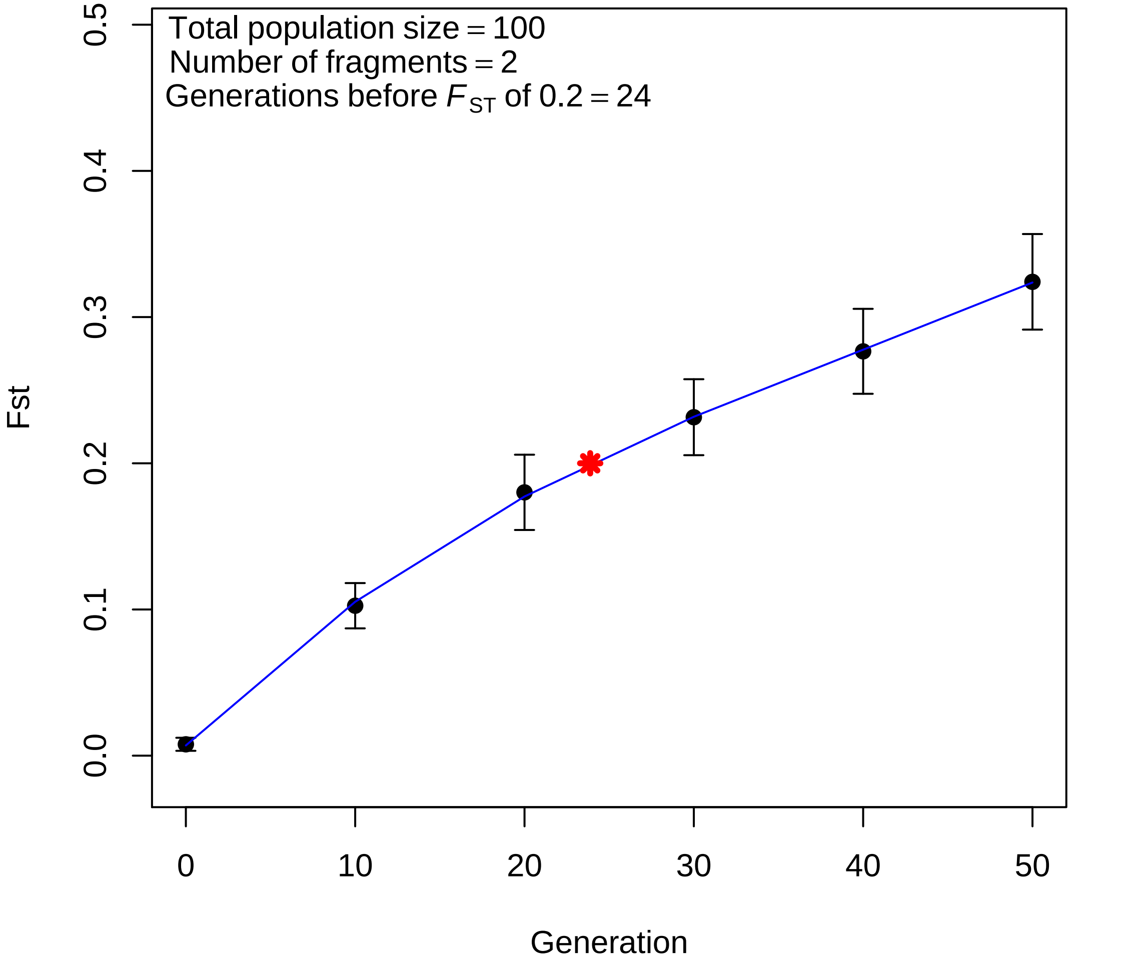

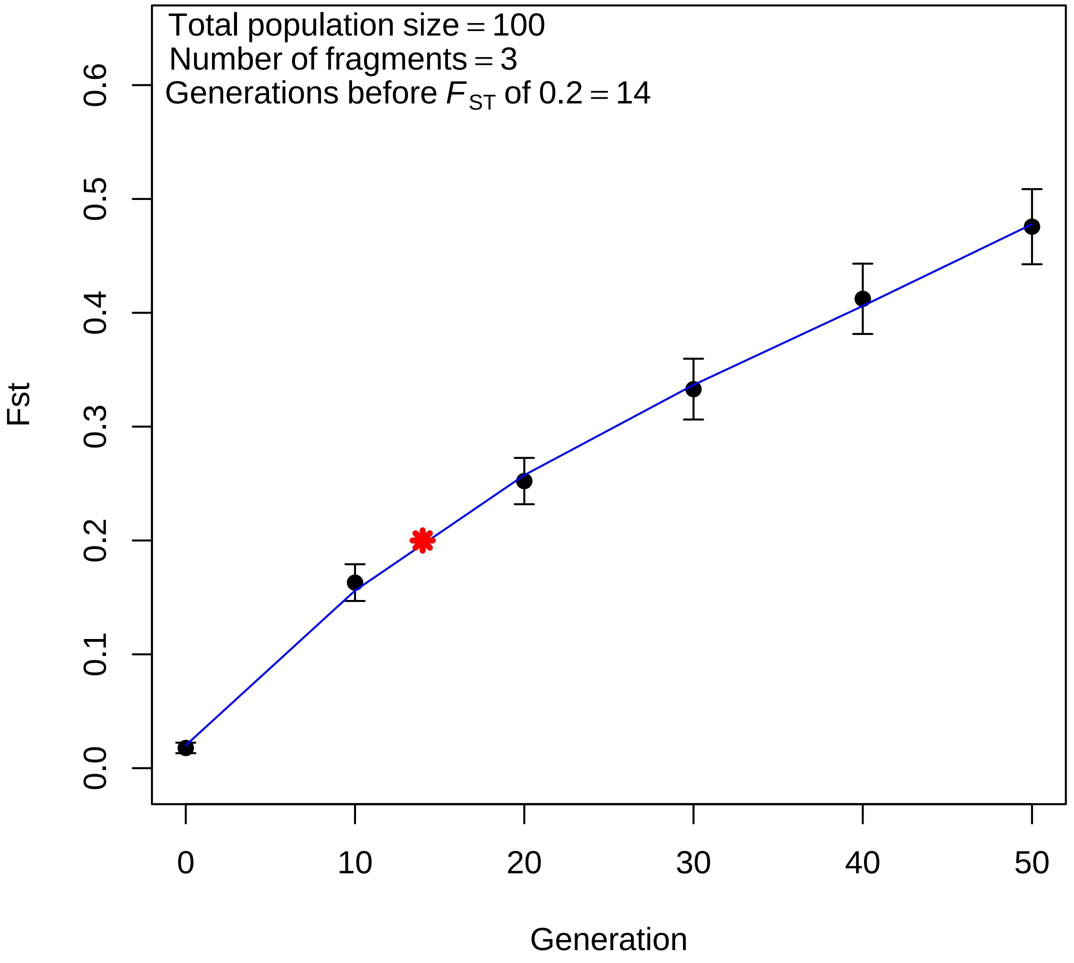

Total population size = 100
Number of fragments = 3
Generations before $F_{ST}$ of 0.2 = 14
Fst
0.0
0.1
0.2
0.3
0.4
0.5
0.6
0
10
20
30
40
50
Generation

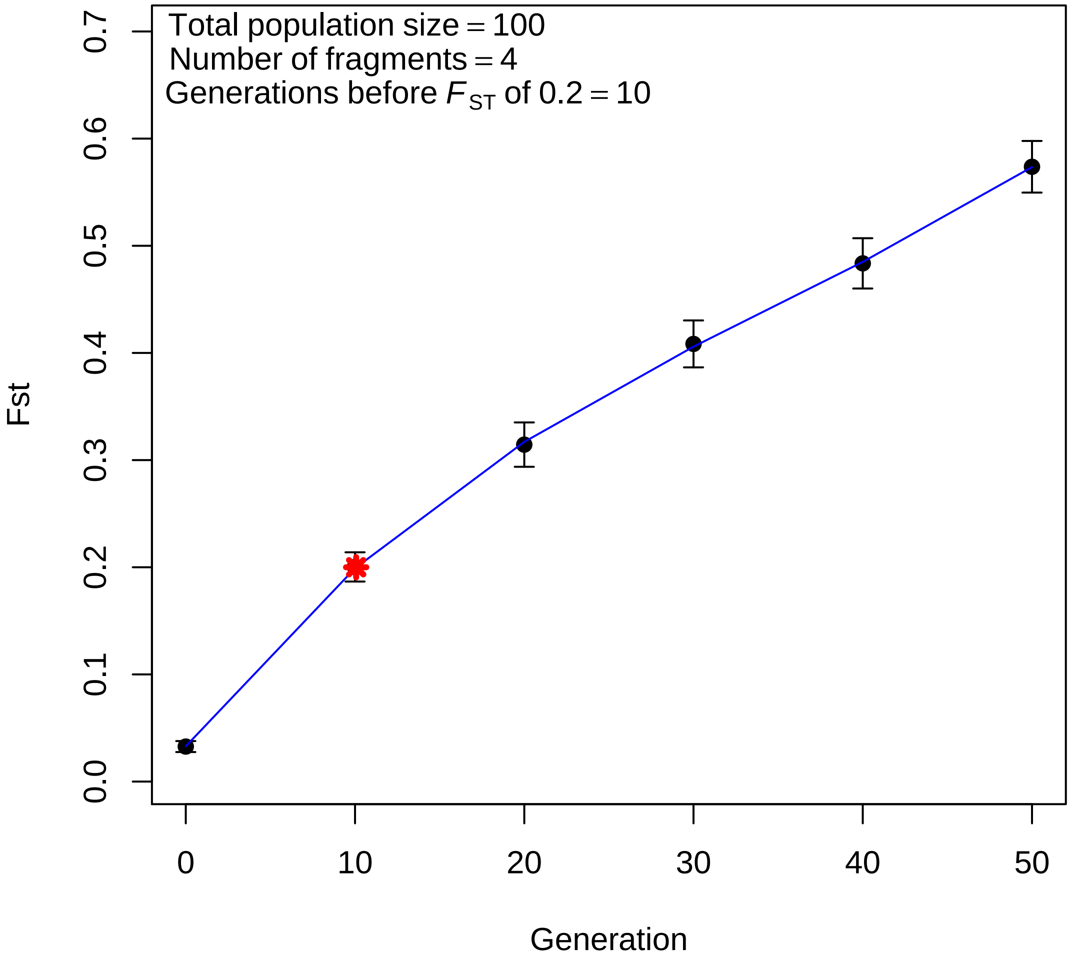
Total population size = 100
Number of fragments = 4
Generations before $F_{ST}$ of 0.2 = 10
Fst
0.0
0.1
0.2
0.3
0.4
0.5
0.6
0.7
0
10
20
30
40
50
Generation

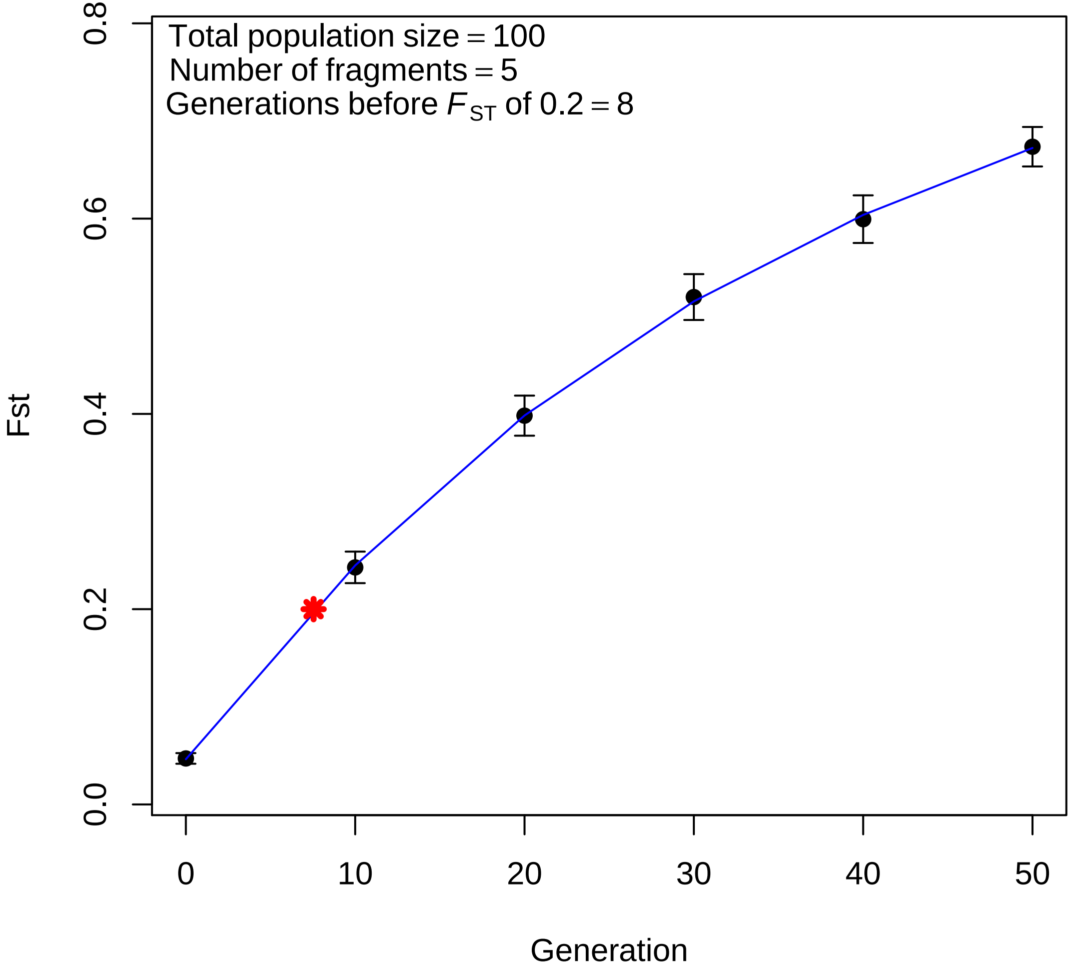
Total population size = 100
Number of fragments = 5
Generations before $F_{ST}$ of 0.2 = 8
Fst
0.0
0.2
0.4
0.6
0.8
0
10
20
30
40
50
Generation

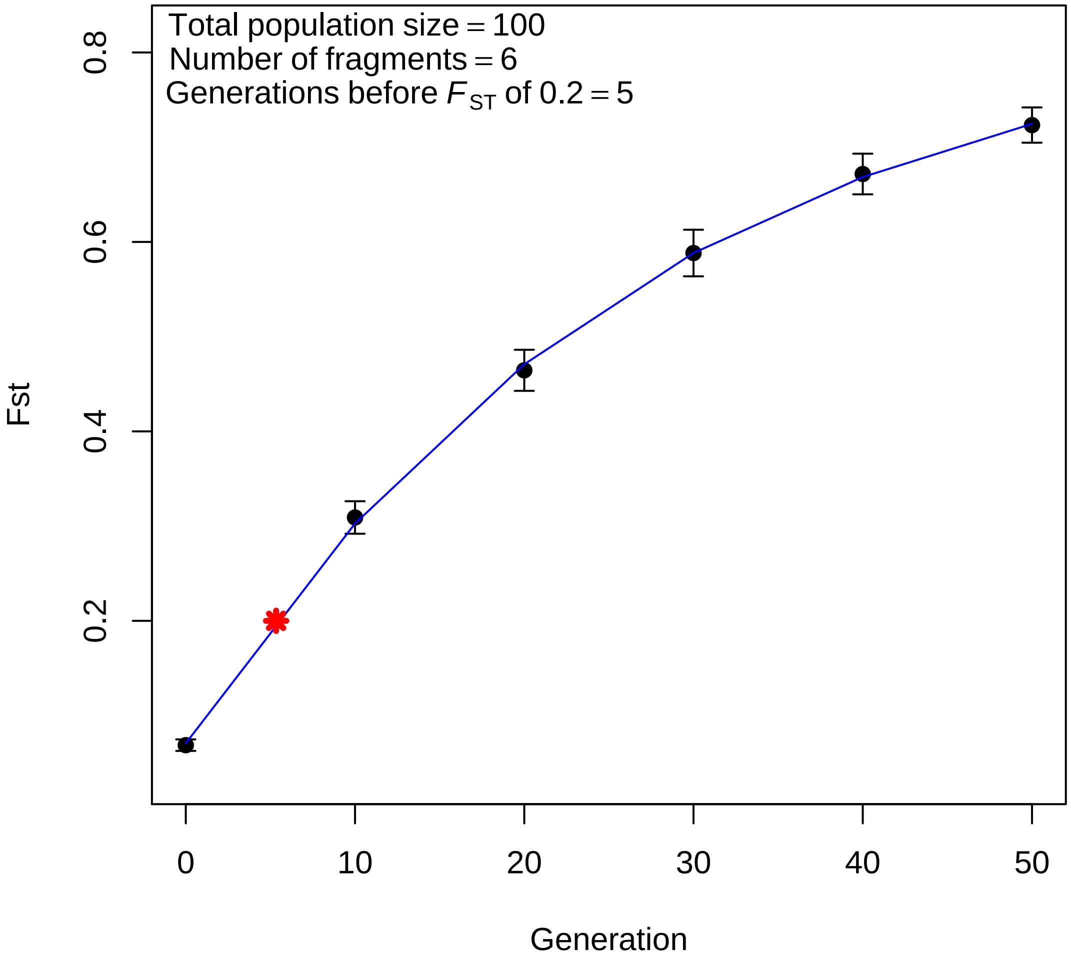

Total population size = 100
Number of fragments = 6
Generations before $F_{ST}$ of 0.2 = 5
Fst
0.2
0.4
0.6
0.8
0
10
20
30
40
50
Generation

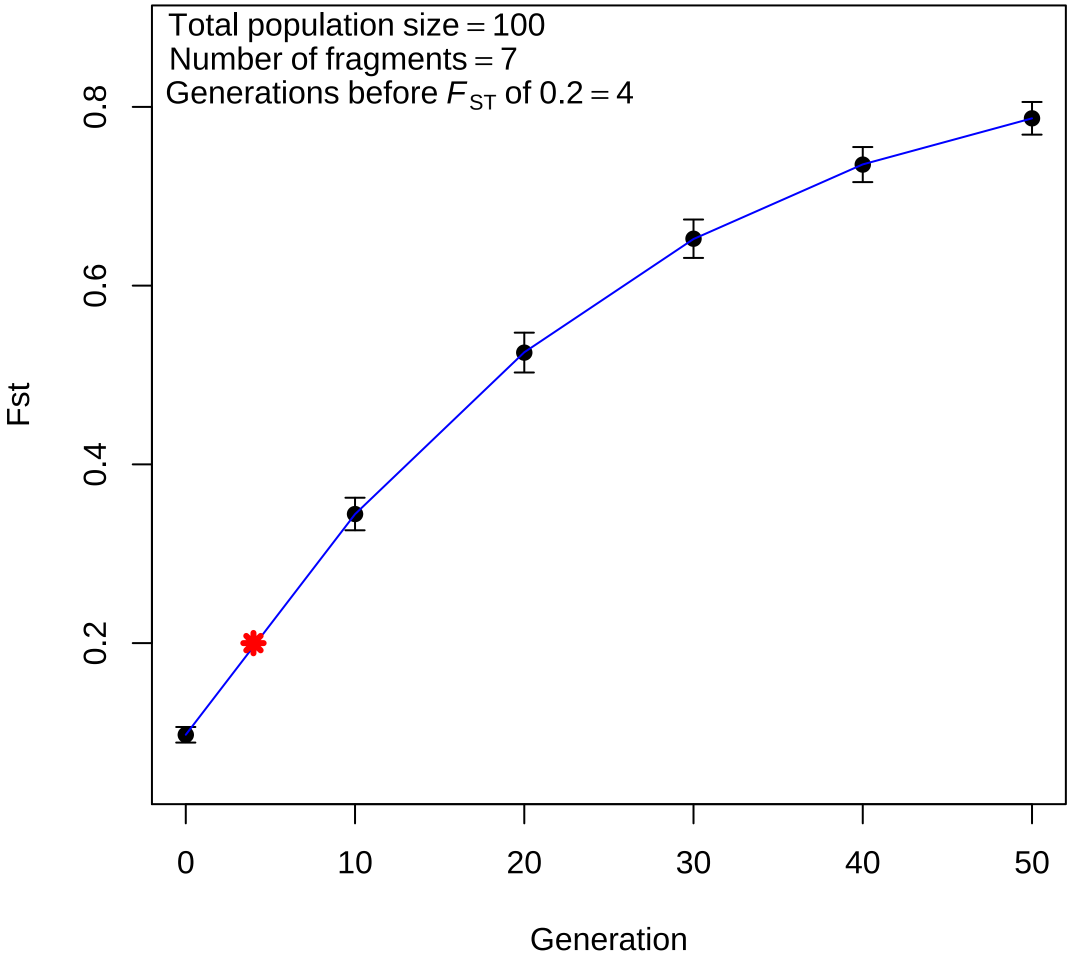

Total population size = 100
Number of fragments = 7
Generations before $F_{ST}$ of 0.2 = 4
Fst
0.2
0.4
0.6
0.8
Generation
0
10
20
30
40
50

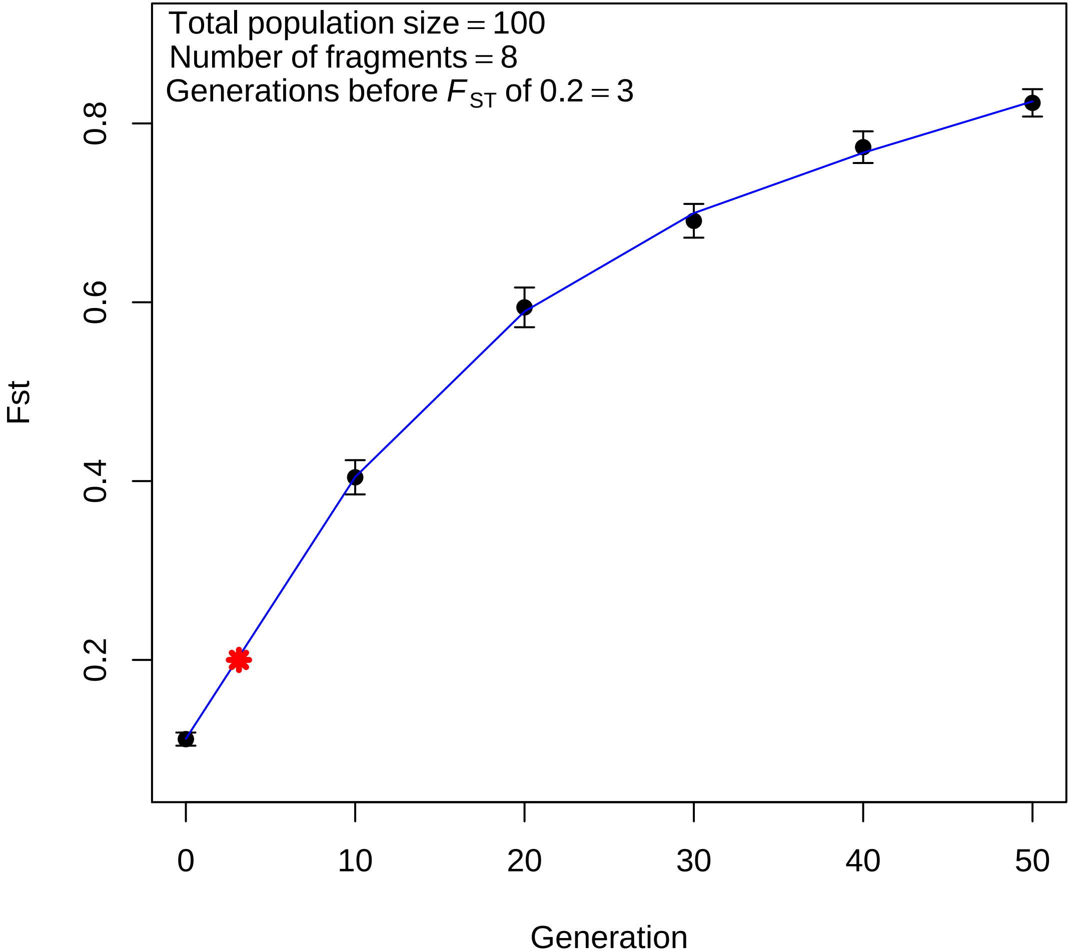

Total population size = 100
Number of fragments = 8
Generations before $F_{ST}$ of 0.2 = 3
Fst
0.2
0.4
0.6
0.8
0
10
20
30
40
50
Generation

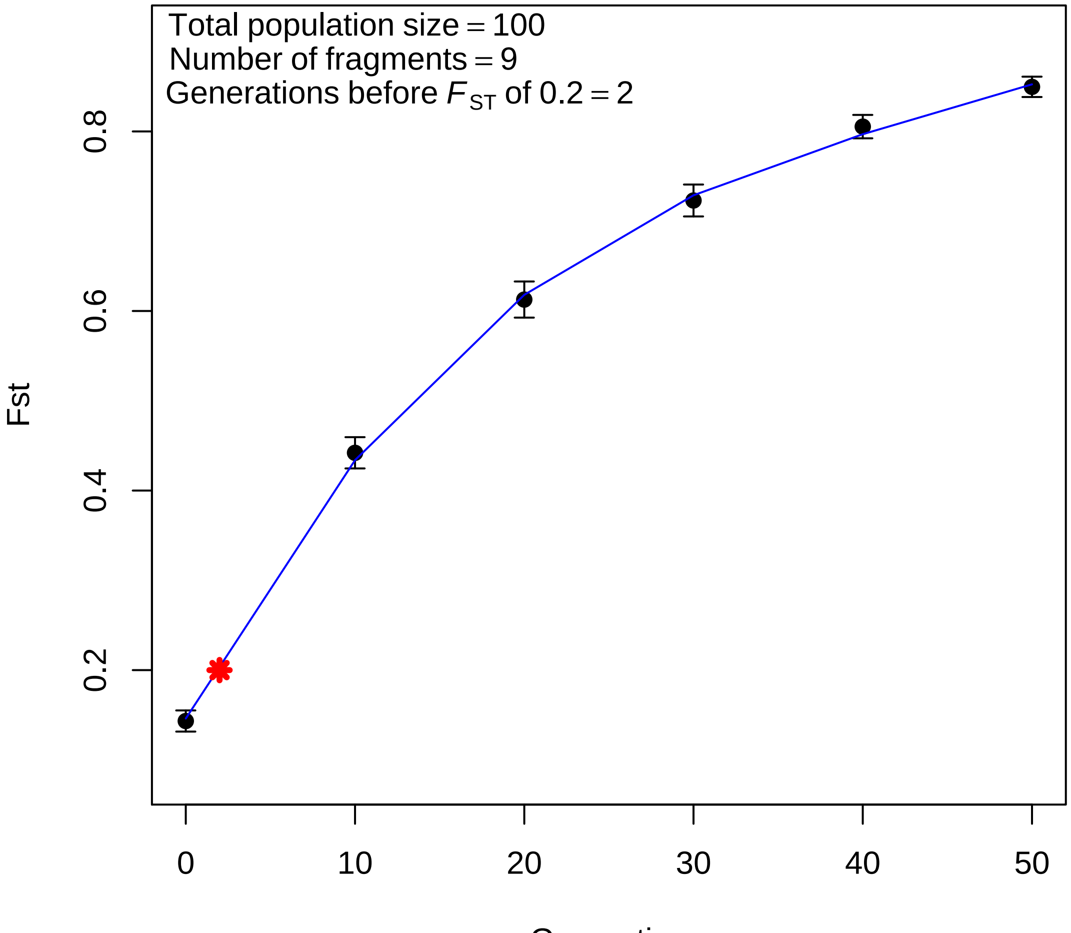
Total population size = 100
Number of fragments = 9
Generations before $F_{ST}$ of 0.2 = 2
Fst
0.2
0.4
0.6
0.8
0
10
20
30
40
50
Generation

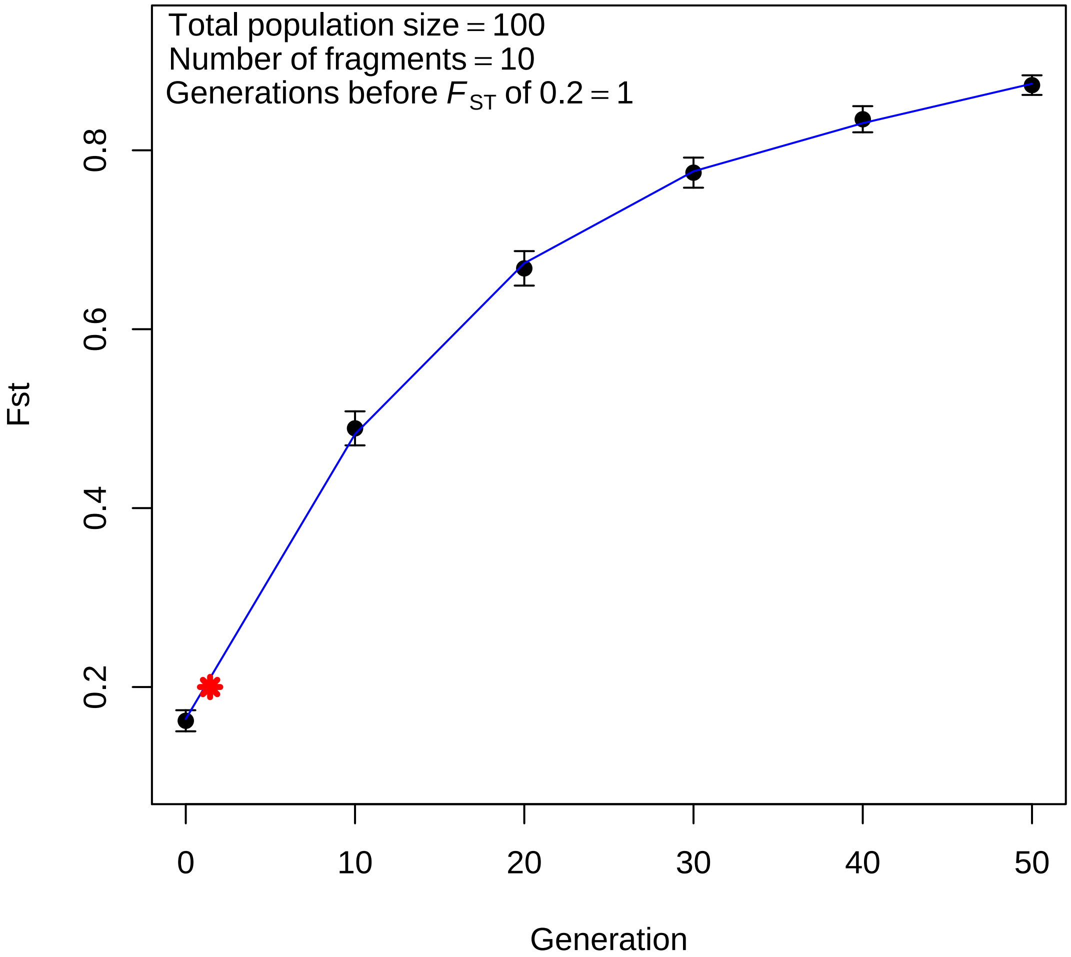

Total population size = 100
Number of fragments = 10
Generations before $F_{ST}$ of 0.2 = 1
Fst
0.2
0.4
0.6
0.8
0
10
20
30
40
50
Generation
